# Aging-associated Increase of GATA4 levels in Articular Cartilage is Linked to Impaired Regenerative Capacity of Chondrocytes and Osteoarthritis

**DOI:** 10.1101/2025.03.18.643933

**Authors:** Meagan J. Makarczyk, Yiqian Zhang, Alyssa Aguglia, Olivia Bartholomew, Sophie Hines, Kate Li, Suyash Sinkar, Silvia Liu, Craig Duvall, Hang Lin

**Author notes:** Correspondence: Hang Lin, PhD, 350 Technology Drive, Room 417, Pittsburgh, PA, 15219, USA. These authors contributed equally (Meagan J. Makarczyk, Yiqian Zhang).

## Abstract

Although the causal association between aging and osteoarthritis (OA) has been documented, our understanding of the underlying mechanism remains incomplete. To define the regulatory molecules governing chondrocyte aging, we performed transcriptomic analysis of young and old human chondrocytes from healthy donors. The data predicted that GATA binding protein 4 (GATA4) may play a key role in mediating the difference between young and old chondrocytes. Results from immunostaining and western blot showed significantly higher GATA4 levels in old human or mouse chondrocytes when compared to young cells. Moreover, overexpressing *GATA4* in young chondrocytes remarkably reduced their cartilage-forming capacity *in vitro* and induced the upregulation of proinflammatory cytokines. Conversely, suppressing *GATA4* expression in old chondrocytes, through either siRNA or a small-molecule inhibitor NSC140905, increased the production of aggrecan and collagen type II, and also decreased levels of matrix-degrading enzymes. In OA mice induced by surgical destabilization of the medial meniscus, intraarticular injection of lentiviral vectors carrying mouse *Gata4* resulted in a higher OA severity, synovial inflammation, and pain level when compared to control vectors. Mechanistically, we found that overexpressing GATA4 significantly increased the phosphorylation of SMAD1/5. Our work demonstrates that the aging-associated increase of GATA4 in chondrocytes plays a vital role in OA progression, which may also serve as a target to reduce osteoarthritis in the older population.

## Introduction

Aging is an inevitable phenomenon resulting in limited functionality, loss of structural integrity, and inability to effectively resist injury and disease.^1^ Osteoarthritis (OA), the world’s most common form of degenerative disease, has been closely associated with the advancement of age.^2^ Specifically, OA is estimated to affect 32.5 million Americans, with most cases occurring in adults over the age of 45.^3^ In fact, 1 in 3 people over the age of 65 are suffering from OA.^4^ There are many contributors linked to the onset and progression of aging, such as organelle dysfunction and telomere shortening,^1^ but the molecular mechanism underlying age-related OA has not been fully understood. Prior studies have demonstrated the role of DNA damage on chondrocyte aging due to oxidative stresses, resulting in an aged phenotype.^5,6^ Aged cells can also undergo a state similar to permanent cell cycle arrest known as cellular senescence, which leads to a low-grade state of chronic inflammation and contributes to OA onset and progression in aged individuals.^6–8^

OA is now considered a whole joint disease, but cartilage degradation still represents the central feature.^3^ The physiological role of articular cartilage is to support and protect the bones of diarthrodial joints through absorbing mechanical loads and facilitating frictionless movements. The extracellular matrix (ECM) of the cartilage is composed primarily of collagen type II and glycosaminoglycans (GAGs), and the constant breakdown and rebuilding of ECM components in the cartilage is common in healthy adults.^3,9^ Many growth factors contribute to the chondro-supportive environment in the knee joint. Particularly, transforming growth factor-β (TGF-β) plays a key role in maintaining chondrocytes and replenishing ECM loss. However, during OA, TGF-β can induce catabolic processes in chondrocytes, resulting in matrix stiffening, osteophytes, and chondrocyte hypertrophy.^10–12^ Furthermore, as we age, the regenerative response of chondrocytes begins to decline.^2^ The whole joint nature of OA and the limited regenerative capacity of chondrocytes have contributed to the difficulty in developing disease-modifying osteoarthritis drugs (DMOADs). To date, no DMOADs have reached FDA approval.^3^

Currently, OA-associated changes in chondrocytes have been widely examined, which have significantly enhanced our understanding of this disease and promoted the development of potential treatments. However, OA is a combined consequence of different physiological stressors, including aging, injuries, obesity, etc. Therefore, aged chondrocytes in healthy humans without OA do not necessarily exhibit all the features of OA chondrocytes. In this study, to understand key regulators that contribute to an “aged” state in chondrocytes prior to OA onset, we first compared the transcriptome of young and old human chondrocytes isolated from healthy donors without joint diseases and employed unbiased analysis to define the key regulatory molecules that mediate chondrocyte aging. Assessment of the chondrocyte genome demonstrated the upregulation of several factors in aged chondrocytes, and transcription factor GATA binding protein 4 (GATA4) specifically drew our attention since it is associated with DNA damage and cellular senescence.^5,13^ Mechanistically, upregulation of GATA4 was shown to increase nuclear factor-κB (NF-κB) pathway activation.^14,15^ NF-κB is thought to amplify and potentially propagate cellular senescence during the aging process through the senescence-associated secretory phenotype (SASP), which could contribute to a low-grade state of chronic inflammation.^16^ Furthermore, the upregulation of GATA4 in OA chondrocytes was also reported.^17,18^ We thus hypothesized that the increased GATA4 level contributes to chondrocyte aging and accelerated OA progression upon injuries. We first tested the hypothesis by increasing or suppressing *GATA4* expression in healthy chondrocytes and examining their cartilage-forming capacity. When *GATA4* was overexpressed, we found that there were alterations to the TGF-β signaling pathway and activation of the NF-κB signaling pathway. We also assessed the role of GATA4 *in vivo* by injecting lentiviral vectors carrying control or mouse *Gata4* genes into the knee joints of OA mice induced by surgically created destabilization of the medial meniscus (DMM). Lastly, a mechanistic study was conducted to explore how GATA4 impacts chondrocyte phenotypes and functions.

## Materials and Methods

### Cell isolation and expansion

Healthy human knee cartilage tissues were harvested from arthritis-free donors through an established protocol with the National Disease Research Interchange (NDRI). This study was approved by the University of Pittsburgh Committee for Oversight of Research and Clinical Training Involving Decedents. Cartilage was diced into ∼ 1 mm^3^ pieces with a scalpel and incubated with a dissociation medium that was composed of high glucose Dulbecco’s Modified Eagle Medium (DMEM, Gibco/Thermo Fisher Scientific, Waltham, MA, United States), 2% Antibiotics-Antimycotics (Life Technologies, Carlsbad, CA, United States), and collagenase type II (1 mg/mL(w/v), Worthington Biochemical Corporation, Lakewood, NJ, United States). 10 mL medium was used for 1g of cartilage, and the treatment lasted for 16 hours in a shaker at 37°C. The mixture was then passed through a 70 µm strainer to collect single chondrocytes. Isolated chondrocytes were seeded in tissue culture flasks at 1×10^4^ cells/cm^2^ and maintained in growth medium (GM, DMEM containing 10% fetal bovine serum (FBS, Life Technologies) and 1% Antibiotics-Antimycotics). After cells were fully attached to the culture substrate, the medium was changed every three days until cells reached 70-80% confluency. Cells were detached with Trypsin/EDTA (Gibco/ Thermo Fisher Scientific) and passaged.

### Individual Chondrocyte Monolayer Culture

Chondrocytes isolated from healthy human cartilage were expanded to passage 1 (P1) and plated in tissue culture 6 well plates at P2, treated with GM, and cultured until cells reached 80% confluency. Media changes occurred every two days until cell collection. **Table S1** lists the chondrocyte ages and genders used for different experiments.

### Young and Old Chondrocyte Pools

Chondrocytes at passage 0 (P0) were pooled with three to four other young chondrocyte donors and grown in tissue culture flasks with GM at a cell seeding density of 1×10^6^ cells per flask. Media was changed weekly. Once cells reached 80% confluency, cells were collected using Trypsin/EDTA. Some cells were frozen using Recovery Cell Culture Freezing Medium (Gibco/ Thermo Fisher Scientific), and other cells were used for subsequent passaging. The same methods were used for the old (> 45 years) chondrocyte pool. **Table S1** lists the chondrocyte ages and genders used in each pool. Due to the extensive amount of work conducted using these pools, two separate chondrocyte pools were made.

### RNA sequencing (RNA-Seq) and bioinformatics analysis

The transcriptomic differences among three young and three old chondrocyte donors were assessed through RNA-Seq. Individual chondrocyte cultures (P0) were lysed with the QIAzol reagent (Qiagen, German Town, MD, United States), and RNA was isolated from the lysate using an RNA Easy Plus Universal Kit (Qiagen). Extracted RNA were quantitated Qubit™ RNA BR Assay Kit (Thermo Fisher Scientific) followed by the RNA quality check using Fragment Analyzer (Agilent Technologies, Santa Clara, CA). For each sample, RNA libraries were prepared from 500ng RNA using the KAPA mRNA HyperPrep Kit (Roche, Indianapolis, IN) according to manufacturer’s protocol, followed by quality check using Fragment Analyzer (Agilent Technologies) and fluorescent quantification on the Infinite F Nano + (Tecan, Männedorf, Switzerland). The libraries were normalized and pooled, and then sequenced using NovaSeq6000 platform (Illumina, San Diego, CA,) to an average of 50M 100PE reads.

Quality control was first applied to raw RNA sequencing reads by tool FastQC. Low-quality reads and adapter sequences were filtered out by the Trimmomatic tool. Surviving reads were then aligned to human reference genome hg38 using STAR aligner, and gene counts were quantified. Differential expression analysis was performed based on gene counts by R package “DESeq2” and DEGs were selected by adjusted p-value < = 0.05 and fold change > = 1.5. These DEGs were then applied to Ingenuity Pathway Analysis (IPA) to detect enriched pathways. This software employs databases of prebuilt pathways with known genes summarized from previous studies, checks the overlap between the DEG list and known pathways and performs statistical tests to determine the enrichment. Significant pathways were defined by p-value < = 0.05. Statistically stringently, FDR = 5% cutoff should be applied to control the false discovery rate. To encourage more gene candidates, this study went by p-value< = 0.05 and fold-change> = 1.5 cutoff. All the tools were run by default parameter settings.

### Immunohistochemistry (IHC) to examine the GATA4 levels in native cartilage tissues

Human cartilage tissues were fixed in 10% buffered formalin (Fisher Chemical, Fair Lawn, NJ) at 4°C overnight and then subjected to a graded ethanol dehydration series, starting from 20% ethanol and progressing to 100% ethanol. Subsequently, they were embedded in Paraplast X-tra (Leica Biosystems Inc. Richmond, IL). The Paraplast-embedded samples were sectioned at a thickness of 6 µm using a rotary Leica microtome (Leica Microsystems Inc., Deerfield, IL, Model RM 2255). Young and old mouse knee joints were gifts from Dr. Ana Mario Cuervo’s lab at Albert Einstein College of Medicine, which otherwise were wastes. After the specimens were fixed and decalcified in a formic acid-based bone decalcifier (StatLab, Mckinney, TX, USA) for a period of two weeks, they were then embedded and sectioned as described above.

For immunohistochemistry (IHC), the formalin-fixed paraffin-embedded sections first underwent antigen retrieval based on different antibodies. Slides were then blocked with 10% goat or horse serum (Abcam, Cambridge, MA) in phosphate buffered saline (PBS, Thermo Fisher Scientific) for 1h, incubated at 4^◦C^ overnight with the primary antibody against GATA4, then incubated with a biotinylated anti-mouse/rabbit immunoglobulin G(IgG) secondary antibody for 1h, with signal detection via DAB substrate kit (Abcam). The Nikon Eclipse E800 upright microscope (Melivile, NY, United States) was used to image the stained sections. Antibody specifications can be found in **Table S2**.

### Western blot to examine the GATA4 levels in human chondrocytes

P1 young and old chondrocytes were used for individual chondrocyte analysis of GATA4. Cells were washed in pre-cooled PBS (Thermo Fisher Scientific) three times. Using the RIPA buffer (Sigma-Aldrich) supplemented with the protease and phosphatase Inhibitor Single-Use Cocktail (Gibco/Thermo Fisher Scientific) and a cell scraper, monolayer culture samples were collected. A pestle was used to homogenize pellets in the RIPA cocktail for pellet culture samples. The protein concentration of the supernatant was determined by the Pierce^™^ BCA Protein Assay Kit (Thermo Scientific). Proteins were fractioned electrophoretically on the NuPAGE 4-12%, Bis-Tris Mini Protein Gel (Gibco/Thermo Fisher Scientific) and then transferred to a polyvinylidene fluoride (PVDF) membrane using the iBlot Dry Blotting System (Invitrogen, Waltham, MA, United States). The membrane was blocked with 3% non-fat milk (Bio-Rad, Hercules, CA, USA), diluted with 1× Tris-buffered saline (TBS, Gibco/ Thermo Fisher Scientific) and 0.1% Tween 20 (Sigma-Aldrich) (TBST) at room temperature for 1.5 h, washed, and incubated with the primary antibody at 4°C overnight on a rotating shaker. The membrane was washed 7 times for 3 min with TBST buffer and incubated with horseradish peroxidase (HRP)-linked secondary antibodies (GE Healthcare Life Sciences, Malborough, MA, United States) for 1.5 h at room temperature. After being washed 5 times with TBST, the membrane was incubated in the chemiluminescence substrate SuperSignal West Dura Extended Duration Substrate (Thermo Fisher Scientific). Images were acquired using the ChemiDocTM Touch Imaging System (Bio-Rad). Images were quantified using ImageJ. Antibody information is included in **Table S2**.

### Overexpression of *GATA4* in young human chondrocytes

P2 young, pooled chondrocytes or young, individual chondrocytes were transduced with the lentiviral vector containing *GATA4* fused with dTomato gene or the control lentivirus carrying EGFP for 10 hours. After that, flasks were rinsed with PBS for 2 times and the medium was replaced by fresh GM. Both vectors were created and packed by VectorBuilder (>10^8^ TU/mL, VectorBuilder, Chicago, IL, United States). To detect the number of cells transduced, cultures were imaged with an EVOS M5000 microscope (Thermo Fisher Scientific) after 72 h of initial transduction. After transduction, western blot, RT-qPCR, and IHC were used to verify the stable expression of GATA4 in cells.

### RNA isolation and quantitative real-time polymerase chain reaction (RT-qPCR)

For pellet culture, samples were first rinsed with PBS twice and a pestle and electric pulverizer were used to crush pellets. Cells were homogenized in Qiazol (Qiagen). Total RNA was isolated and purified using the RNAeasy Plus Universal Mini Kit (Qiagen, Cat. NO. 74104) according to the manufacturer’s protocol. The reverse transcription to the complementary DNA (cDNA) was accomplished using the SuperScript IV VILO Master Mix (Invitrogen). RT-qPCR was performed on a real-time PCR instrument (QuantStudio 3, Applied Biosystems, Foster City, CA, United States) using the SYBR Green Reaction Mix (Applied Biosystems) with custom primers ordered from Integrated DNA Technologies (IDT, Newark, NJ, United States). Relative gene expression levels were calculated through the 2 ^-ΔΔCt^ method. Ribosomal protein L13A (*RPL13A*) was used as the housekeeping gene. Full names and abbreviations of genes and their corresponding proteins are listed in **Table S3,** and primer sequences are listed in **Table S4**.

### Pellet culture and chondrogenesis of young human Chondrocytes overexpressing *GATA4*

Following the lentiviral transduction of *GATA4* gene in young, pooled chondrocytes, the cells were collected and formed into pellets at a cell seeding density of 3 × 10^5^ cells per pellet. Pellets were treated with chondrogenic medium (CM, DMEM with 1% v/v Insulin-Transferrin-Selenium-Ethanolamine (ITS, Gibco/ Thermo Fisher Scientific), 1% antibiotic-antimycotics, 10^-7^ M dexamethasone (Sigma-Aldrich), 40 µg/mL L-proline (Sigma-Aldrich), supplemented with 10 ng/mL transforming growth factor-β3 (TGF-β3, Peprotech, Rocky Hill, NJ, United States), and 50 µg/mL ascorbic acid-2-phosphate (Sigma-Aldrich)). Medium was changed daily for seven days. RT-qPCR, histology, IHC, and western blot were used to characterize the tissues.

### Safranin O/Fast green staining

Pellet samples were fixed in 10% buffered formalin (Fisher Chemical) for 2 h at room temperature and then rinsed with PBS. Pellets then underwent serial dehydration in 30,50,70,95, and 100% ethanol for one hour each. The 100% ethanol was refreshed once for an additional hour prior to sample clearing in Xylene. Pellets were cleared in Xylene (Fisher Chemical) for two hours. Pellets were then placed in Paraplast X-tra (Leica Biosystems Inc.) overnight. The next day, pellets were embedded in Paraplast X-tra blocks and sectioned at a 6-μm thickness using a Leica microtome (Leica Microsystems Inc., Model RM 2255).

Slides were stained using Safranin O (0.5%, Catalog number: 50240, Sigma-Aldrich), in 1% acetic acid (Catalog number: A6283, Sigma-Aldrich), 0.005% fast green (0.05 g, Catalog number: 104022, Sigma-Aldrich) in 100 mL distilled water (Invitrogen) and counterstained with Hematoxylin QS solution (Catalog number: H3404, Vector Laboratories INC). Imaging was conducted using a Nikon Eclipse E800 upright microscope.

### LUMINEX multiplex assays

Upon 7 days of chondrogenesis, condition medium from pellets was collected and flash frozen in liquid nitrogen and immediately stored in -80°^C^. LUMINEX assays were accomplished using the Bio-Plex 200 system (Bio-Rad). Data collection and analysis were conducted using the Bio-Plex Manager 6.1 software as established in previous studies.^19^ LUMINEX kit information can be found in **Table S5**.

### Knockdown of *GATA4* in old human chondrocytes

The siRNA targeting human *GATA4* (ON-TARGET plus Human GATA4 (2626) siRNA, J-008244-06-0005, Horizon Discovery Biosciences Limited, Cambridge, UK) was used in this study with a scrambled siRNA (ON-TARGET plus non-targeting siRNA #1, Catalog number: D-001810-01-05, Horizon Discovery Biosciences Limited) as the control. Old, pooled chondrocytes were transfected with the siRNA using Lipofectamine RNAiMAX reagent (Thermo Fisher Scientific). After 24 hours of incubation, the transfection medium (Opti-MEM Reduced Serum Medium, Thermo Fisher Scientific) was changed to GM. Transfected cells were collected after 48 hours for RT-qPCR to confirm the knockdown efficiency.

### Pellet culture and chondrogenesis of old human chondrocytes with *GATA4* Knockdown

Following the 48h transfection of scrambled control siRNA or siRNA targeting *GATA4*, cells were collected using Trypsin/EDTA (Thermo Fisher Scientific) and pellets were made at a cell density of 3×10^5^ cells per pellet. Pellets were treated with CM supplemented with 10 ng/mL TGF-β3 and 50 µg/mL ascorbic acid-2-phosphate for 7 days. Upon day 7, pellets were collected for RT-qPCR, western blot, and IHC.

### GATA4 Small Molecule NSC140905

GATA4 small molecule, NSC140905, also known as HCA 42027( Biosynth Ltd, Compton, United Kingdom), was reconstituted to 7.5 mM stock solution using UltraPure™ DNase/RNase-Free Distilled Water (Invitrogen) on a shaker at 37° until completely dissolved. Old, pooled chondrocytes were pelleted and treated with CM supplemented with 10 ng/mL TGF-β3 and 50 µg/mL ascorbic acid-2-phosphate with 100 μM NSC140905 for 14 days. Pellets were collected for IHC, western blot, and RT-qPCR.

### Animal model

All animal experiments were approved by the University of Pittsburgh Institutional Animal Care and Use Committee (IACUC). Young (8 weeks) male C57BL/6 mice were purchased from Jackson Laboratory (Bar Harbor, ME, USA) and maintained in pathogen-free conditions, with no more than five mice per cage. Mice were provided ad libitum access to food and water, and a 12-hour light/dark cycle to simulate natural circadian rhythms. To minimize bias, mice were randomly assigned to either control or GATA4 groups, with 8 mice in each group.

### Intraarticular injection

Intraarticular injections were administered to mice between 10-12 weeks of age under general anesthesia to safeguard the well-being of the animals and to minimize procedural discomfort. Under general anesthesia with 2% isoflurane in an oxygen mixture, the mice were placed in a supine position, and the right knee joint was positioned at a 90-degree flexion to facilitate accurate injection into the joint space. The injection site was meticulously identified medial to the patellar tendon. Using a 29-gauge needle, a volume of 10 µL of lentiviral particles encoding either GATA4 or a control vector (at a concentration >10^8^ TU/mL, VectorBuilder) was precisely administered into the intra-articular space of the right knee. The precision of the injection was ensured by employing a consistent technique across all animals, thereby reducing variability in the delivery of the viral vectors.

### Destabilization of the medial meniscus (DMM) surgery

One week after viral vector injection, DMM surgery was performed to induce the OA model on mice at 11-13 weeks of age.^20,21^ Briefly, a medial parapatellar incision was made to expose the right knee joint, followed by a careful opening of the joint capsule. The anterior medial menisco-tibial ligament was identified and transected using microscissors. The joint capsule and skin were subsequently sutured with 6-0 silk thread. For experimental controls, a sham operation was performed in which the joint capsule was exposed as in the DMM surgery, but the medial menisco-tibial ligament was left intact.

### Knee hyperalgesia

We expected accelerated OA development after *Gata4* overexpression. To observe these differences, we used six weeks as the time point at which control mice had begun displaying mild OA symptoms. Knee hyperalgesia was evaluated using a Pressure Application Measurement (PAM) device (Ugo Basile, Varese, Italy).^22,23^ Mechanical stimuli were applied to the mouse’s knee joint to assess the degree of hyperalgesia based on the applied pressure and the mouse’s response. Six weeks after either DMM or sham surgery, the mice were carefully removed from their cages to ensure they remained calm and unstimulated. The mouse was held securely by its back to maintain a straight posture, with the right knee joint flexed at approximately 90 degrees. The PAM device was placed on the index finger of the right hand of testers, which, along with the thumb, was used to apply pressure to the medial side of the knee joint, ensuring proper contact. Pressure was gradually applied at a constant rate of 30 g/s while monitoring the pressure curve displayed on the computer. The sensor was released immediately when the mouse exhibited a response to the applied stress, such as head movement, vocalization, or knee withdrawal. The pressure value displayed by the software, and the maximum pressures that mice can withstand were recorded. Two measurements were taken per knee, one on the medial side, and one on the lateral side the average was calculated for accuracy.

### Synovial inflammation score

After 6 weeks, knee joints were collected and sectioned as described above. Histology staining and IHC were used to assess OA severity. Synovial inflammation score was evaluated according to a scoring protocol outlined in previous studies^24^, which relies on the hematoxylin and eosin (H&E)-stained tissue sections.

### Statistical Analysis

Each experiment was carried out with at least three biological replicates. Data is presented as mean ± standard deviation unless otherwise specified. Detailed information of sample size, pre processing, and statistical methods have been specified in each figure legend. Prism 10 (GraphPad, San Diego, CA) was used for statistical analysis. The significance level was set at 0.05 and indicated by * (p < 0.05), ** (p < 0.01), *** (p < 0.001), and **** (p < 0.0001).

## Results

### GATA4 is predicted to regulate chondrocyte aging

To limit the influence of *in vitro* expansion on cell phenotype, P0 human chondrocytes were used for RNA sequencing (**Figure 1A**). The volcano plot showed that 303 genes are upregulated and 163 genes are downregulated in aged chondrocytes compared to young cells (**Figure 1B**). The top 50 most changed genes are listed in **Figure 1C**. Interestingly, the osteoarthritis pathway was identified to be activated in old chondrocytes (**Figure 1E**).

**Figure 1:**
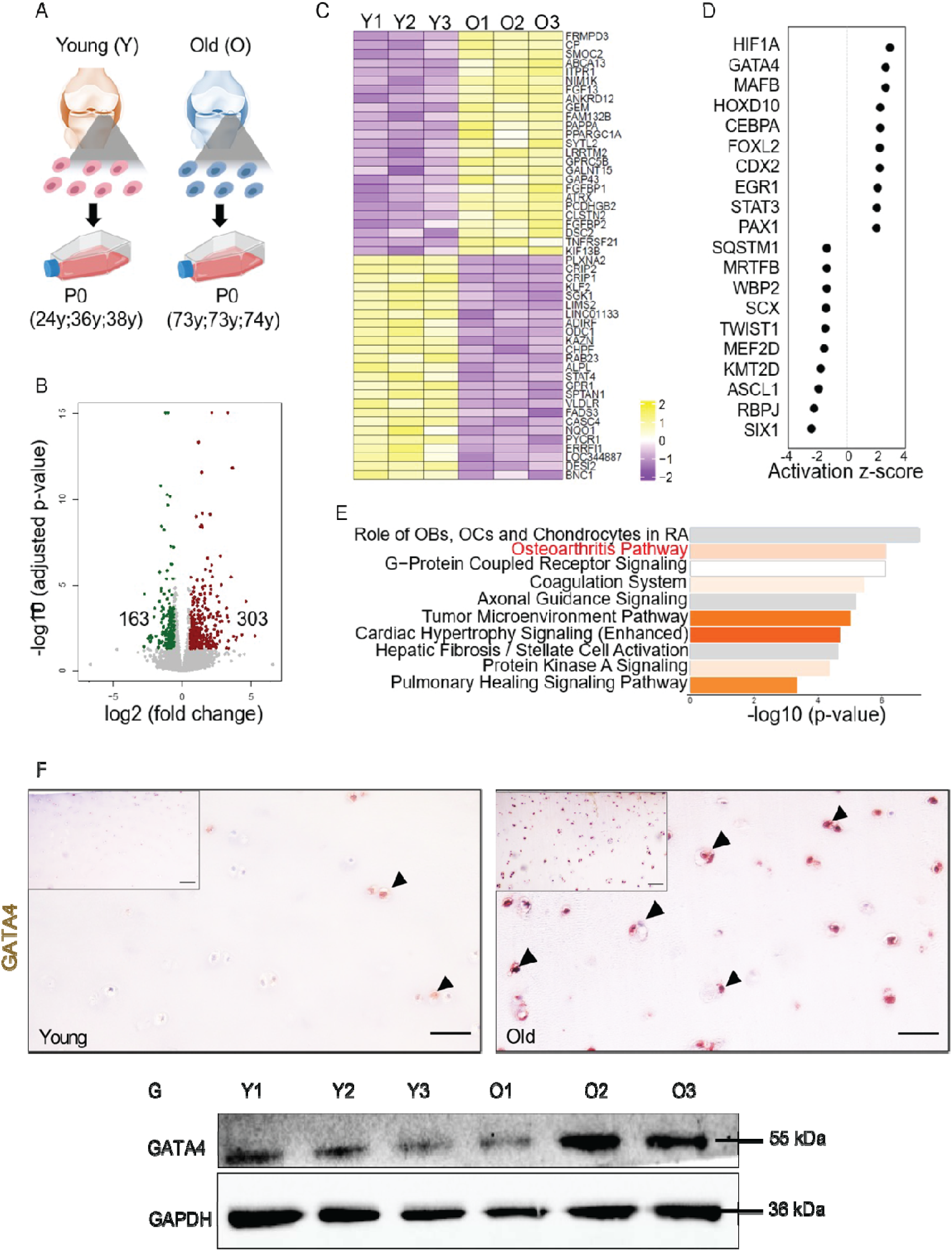
GATA4 is predicted to regulate chondrocyte aging. (A) Healthy chondrocytes were isolated from knee joint cartilage from young and old donors without osteoarthritis (assessed by experienced surgeons). P0 cells were used for RNA sequencing analysis. (B) Volcano plot demonstrating 303 upregulated and 163 downregulated genes in old chondrocytes when compared to young cells. (C) Top 50 genes that are significantly differently expressed in young and old chondrocytes. (D) Activation Z score of top 10 transcription regulators that are activated (positive) or inhibited (negative) in aged versus young chondrocytes. A comprehensive list of gene names can be found in **Table S3**. (E) Ingenuity Pathway Analysis (IPA) of young versus old cells. The gray bars indicate that no activity pattern is identified in IPA despite the highly significant association of the genes within the pathway. Orange, positive z-score; white, zero z-score. OBs=osteoblasts; OCs=osteoclasts. (F) GATA4 IHC of healthy human cartilage tissue from young and aged donors. Bar=50 µm. (G) Relative protein levels of GATA4 in P1 chondrocytes from individual human donors were analyzed by western blot.

Through Ingenuity Pathway Analysis (IPA) Upstream Regulator Analysis, transcriptional regulators that may mediate the difference between young and old cells were predicted, and the top 20 of them are shown in **Figure 1D**. Given that GATA4 was previously shown to regulate chondrocyte senescence^13^ and be involved in the activation of Nuclear Factor Kappa B (NF-κB) pathway in tissues like the nucleus pulposus^25^ and synovium,^26,27^ it was selected for further investigation in this study. It is important to clarify that selecting GATA4 in the current study does not diminish the significance of other molecules involved in chondrocyte aging. Notably, hypoxia-inducible factor 1α (HIF1A) was predicted to be the most active transcription regulator. All the molecules depicted in Figure 1D can be validated and examined in future studies.

Initial data analysis predicted that GATA4 was upregulated in aged chondrocytes compared to young donors (**Figure 1D**). Immunohistochemistry (IHC) results indicated that GATA4 was more abundant in articular cartilage harvested from aged individuals (**Figure 1F**) (**Supplementary Figure S1**). Analyzing knee joints collected from young and old mice also demonstrated that GATA4 levels were higher in aged cartilage tissues compared to young (**Supplementary Figure S2**). To further confirm the findings, western blot was used to examine GATA4 levels in isolated chondrocytes, and the results indicated that aging correlates with the increase of GATA4 in chondrocytes (**Figure 1G**).

#### Overexpressing GATA4 impairs the hyaline cartilage formation capacity of young chondrocytes

To examine the functions of GATA4 in chondrocytes, we first overexpressed it in healthy young human chondrocytes (**Figure 2A**). Real-time quantitative PCR (RT-qPCR), immunostaining, and western blot confirmed the significantly increased GATA4 expression after infection, which did not impact the expression levels of collagen type II (COLII)-α1 (*COL2A1*) and aggrecan (*ACAN*), but significantly upregulated the expression of Collagen type X (COLX)-α1 (*COL10A1*) and India hedgehog (IHH), two representative chondrocytic hypertrophy markers (**Supplementary Figure S3**). We then examined the cartilage formation capacity of cells by culturing cells in chondrogenic medium for 7 days. IHC, RT-qPCR, and western blot demonstrated the continuous overexpression of *GATA4* in newly formed tissues (**Figure 2B-D**). Interestingly, significantly reduced GAG and COLII production was found in the overexpression *GATA4* group (GATA4 group) compared to the GFP control group (**Figure 2E-G**). Similar results were observed in the study testing chondrocytes from individual donors (**Supplementary Figure S4**). Interestingly, *GATA4* overexpression did not impact the expression of *COL10A1* and *IHH* in the cartilage pellets **(Figure 2G**). Additionally, tissues from the GATA4 group secreted more proinflammatory cytokines, including IL-6, IL-8, TNF-α, and chemokine CCL2 (**Figure 2H&I**), as well as enzymes that can break down cartilage, including matrix metalloproteinases *(MMP)-1, 2, 12*, and a disintegrin and metalloproteinases *(ADAMTS) 5* (**Figure 2J&K**). Although the expression levels of *MMP-13* gene decreased in GATA4 group, the protein levels showed no difference between the two groups (**Supplementary Figure S5**).

**Figure 2:**
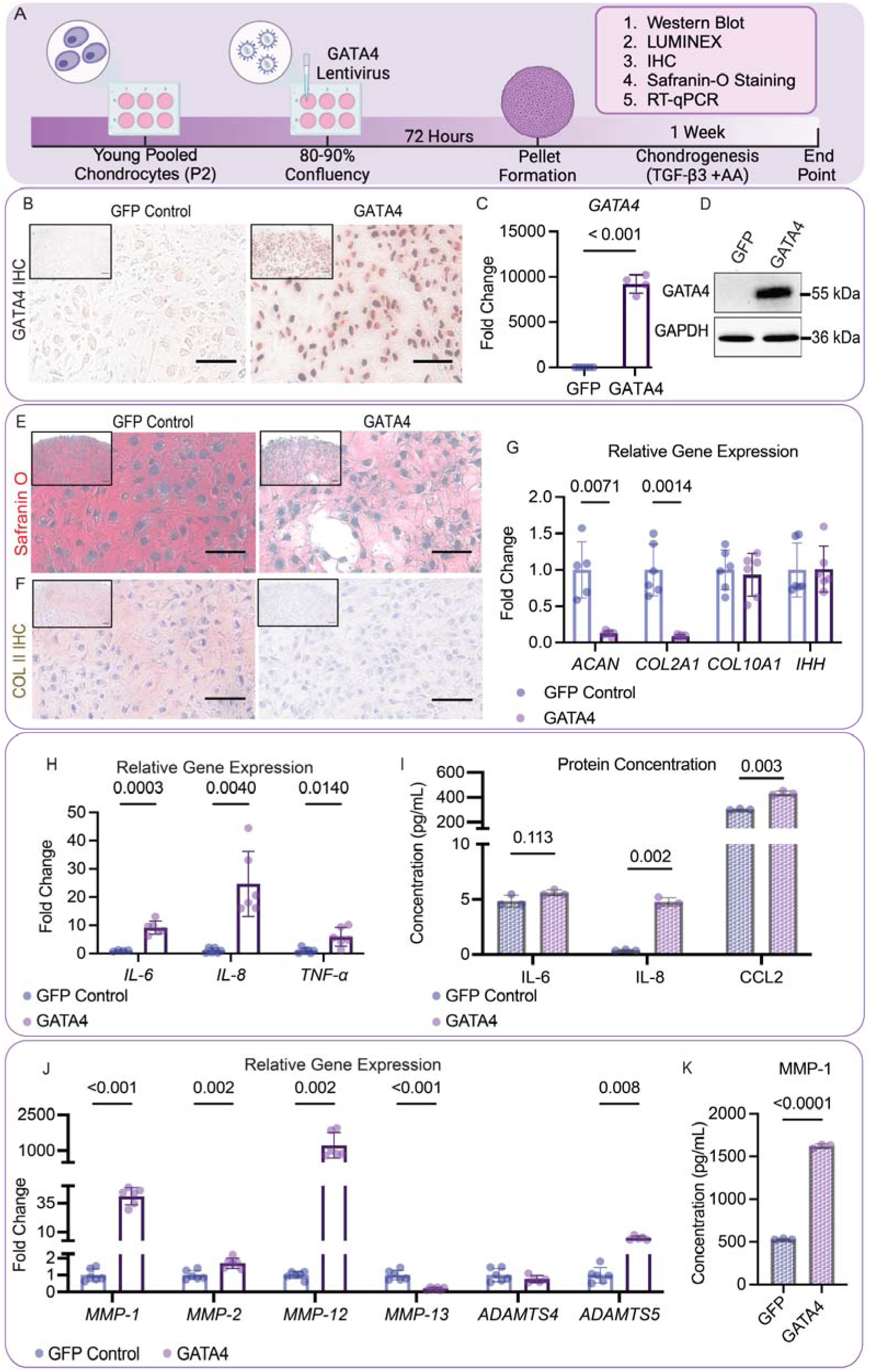
Overexpressing GATA4 impairs the hyaline cartilage formation capacity of young chondrocytes. (A) Timeline depicting the study. (B) IHC to assess GATA4 protein levels in young cells overexpressing GFP control or GATA4. Scale Bar=50 µm. (C) RT-qPCR analysis of GATA4 gene expression in two groups. (D) Western blot to measure GATA4 protein levels. (E) Safranin O staining and (F) Collagen type II (COLII) IHC to examine the production of cartilage matrix. Scale Bar=50 µm. (G) RT-qPCR analysis of gene expression of cartilage matrix proteins aggrecan (ACAN) and collagen type II-α1 (COL2A1) and hypertrophy markers collagen type X-α1 (COL10A1) and India hedgehog (IHH). (H) RT-qPCR analysis of gene expression of proinflammatory cytokines, including interleukin (IL)-6, IL-8, and tumor necrosis factor-α (TNF-α) (n=6). (I) Concentrations of IL-6, IL-8, and chemokine (C-C motif) ligand 2 (CCL2) in condition medium (n=3). (J) RT-qPCR analysis of relative gene expression of matrix-degrading enzymes, including matrix metalloproteinases (MMP)-1,2,3,12, and 13, and a disintegrin and metalloproteinase (ADAMTS) 4 and 5 (n=6). (K) MMP-1 concentration in condition medium (n=3). Student’s two-tailed t-test with Welch’s correction for standard deviation and a p-value of 0.05 was used for all statistical analysis. Created in BioRender. Makarczyk, M. (2025) https://BioRender.com/luon9t1

#### GATA4 overexpression activates SMAD1/5

Increased SMAD1/5 phosphorylation represents a key feature of aged chondrocytes ^28^. We thus examined whether increased GATA4 levels are associated with SMAD1/5 activation (**Figure 3A**). In the experiment testing chondrocytes from individual donors (**Figure 3B, D-G**), overexpression of GATA4, even without the stimulation of TGF-β3, was sufficient to activate SMAD1/5, but not SMAD2/3. In addition, the group that was co-treated with GATA4 overexpression and TGF-β3 displayed the highest phosphorylated SMAD1/5 (pSMAD1/5) levels in all tested groups. Of note, activation of SMAD2/3 was not impacted by GATA levels. A similar trend was also observed in the study using pooled chondrocytes (**Figure 3C**).

**Figure 3.**
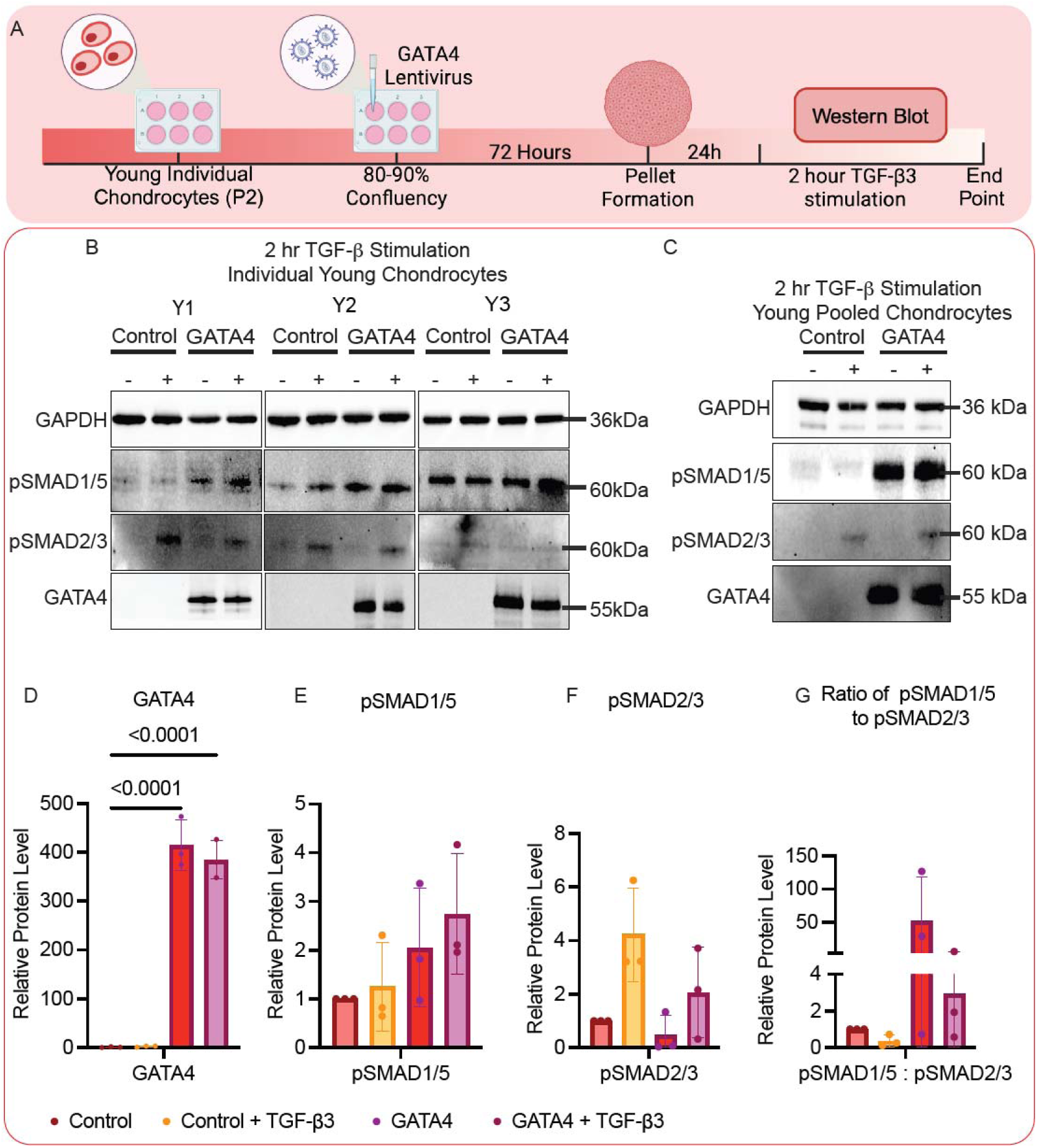
GATA4 overexpression activates SMAD1/5. (A) Schematic showing the experiment. (B, C) Western blot to assess protein levels of phosphorylated SMAD1/5 (pSMAD1/5), phosphorylated SMAD2/3 (pSMAD2/3), and GATA4 in pellets derived from young (B)individual(Y1-3) or (C) pooled chondrocytes, which were infected with lentiviral vectors carrying GATA4 or control genes and then stimulated with (+) or without (-) TGF-β3 for two hours. Relative protein levels of (D)GATA4, (E)pSMAD1/5, and (F)pSMAD2/3 were semi-quantified using ImageJ (n=3). (G) The ratio of pSMAD1/5 compared to pSMAD2/3 was also calculated. Statistics were conducted using one-way Analysis of Variance (ANOVA) with Dunnett’s post hoc analysis. Created in BioRender. Makarczyk, M. (2025) https://BioRender.com/luon9t1

#### Suppressing GATA4 in old chondrocytes promotes ECM formation and lowers proinflammatory cytokines

We then tested the potential of suppressing GATA4 in reversing chondrocyte aging (**Figure 4A**). Several GATA4 siRNAs were tested to examine their capacity to suppress *GATA4*. Based on RT-qPCR results, siRNA2 was selected to be used in all following studies because it induced the lowest expression of *GATA4*. (**Supplementary Figure S6**). *GATA4* knockdown resulted in increased cartilage formation from old chondrocytes, which did not influence the expression of hypertrophy marker *COL10A1* (**Figure 4B-D**). Moreover, although we did not see a difference between the scrambled control and GATA4 siRNA groups regarding the expression of *IL-6* and *IL-8*, the protein level of IL-8 was higher in the GATA4 group. Interestingly, the protein level of CCL2 was significantly decreased after GATA4 knockdown (**Figure 4E, F**). We also tested the expression levels of MMPs and ADAMTSs (**Figure 4H-I**). In general, suppressing GATA4 either decreased or caused no significant changes to the levels of these enzymes. In particular, MMP-13 levels were reduced, in both gene and protein levels (in condition medium), after GATA4 knockdown. Mechanistically, GATA4 siRNA treatment also lowered the phosphorylation of SMAD1/5 and p-P65 (**Figure 4G**).

**Figure 4:**
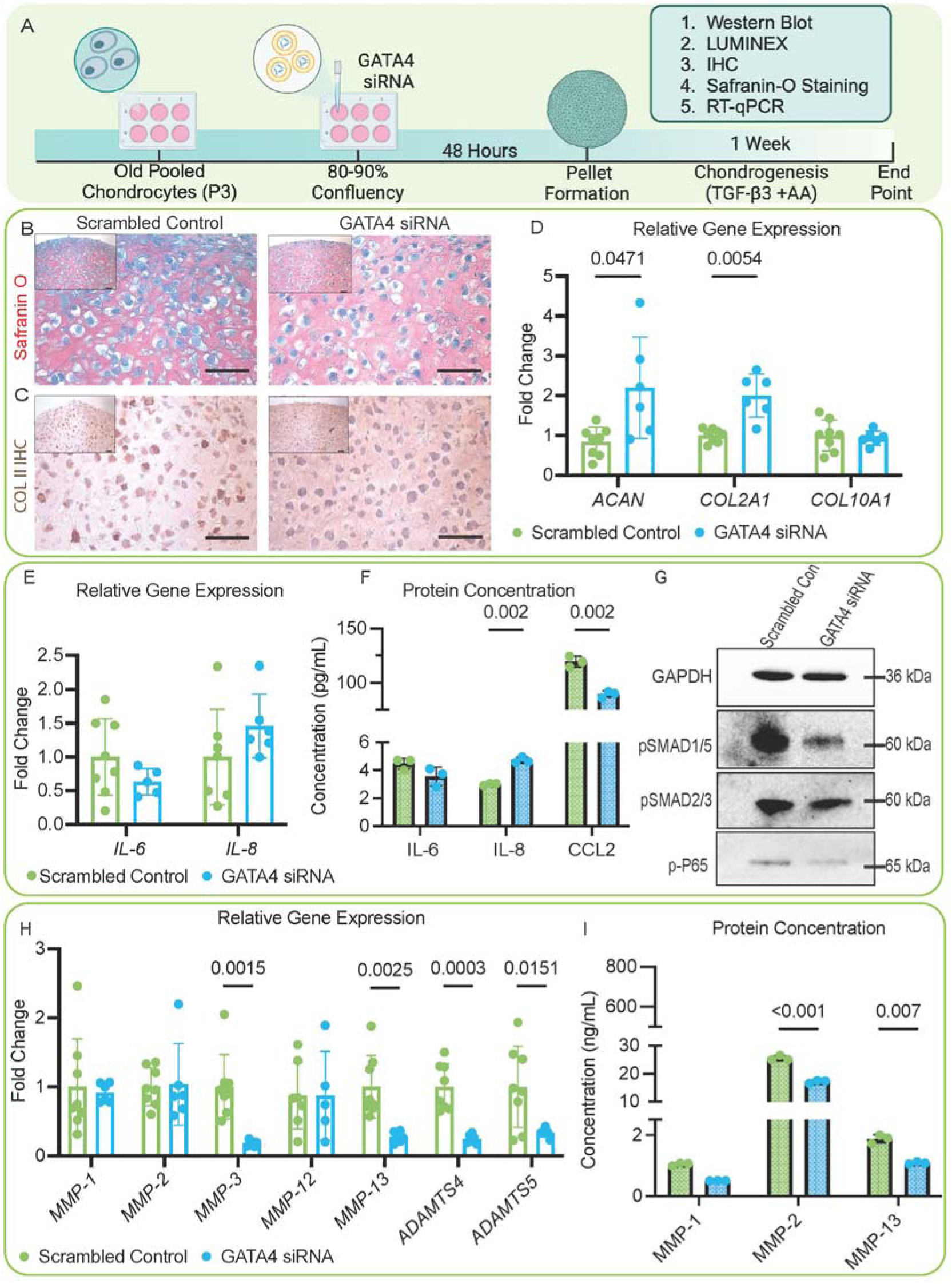
Influence of GATA4 knockdown on in vitro cartilage formation of old chondrocytes. (A) Schematic showing the study. (B) Safranin O staining and (C) COLII IHC to examine the production of cartilage matrix in the Scrambled Control or GATA4 siRNA group. Scale Bar=50 µm. (D) RT-qPCR analysis of relative gene expression of cartilage matrix proteins ACAN and COL2A1 and hypertrophy marker COL10A1(n=6). (E) RT-qPCR analysis of relative gene expression of proinflammatory cytokines IL-6 and IL-8 (n=6). (F) Concentrations of IL-6, IL-8, and chemokine (CCL2) in condition medium (n=3). (G) The relative protein levels of pSMAD1/5, pSMAD2/3, and phosphorylated p65 (p-P65) in two groups. (H) RT-qPCR analysis of relative gene expression of matrix-degrading enzymes, including MMP-1,2,3,12, and 13, and ADAMTS 4 and 5 (n=6). (I) MMP-1,2 and 13 concentrations in condition medium (n=3). Student’s two-tailed t-test with Welch’s correction for standard deviation and a p-value of 0.05 was used for all statistical analysis. Created in BioRender. Makarczyk, M. (2025) https://BioRender.com/luon9t1

In a separate study, we used a small-molecule GATA4 inhibitor NSC140905, which was shown to significantly promote cartilage formation from old chondrocytes and reduce the expression of proinflammatory cytokines (**Supplementary Figure S7**). Taken together, suppressing GATA4 partially restored the capacity of old chondrocytes to create new cartilage.

#### Gata4 overexpression in the knee joints accelerates OA progression in mice

The physiological functions of GATA4 were further examined using a mouse model (**Figure 5A**). Specifically, lentiviral vectors carrying *Gata4* or *mCherry* genes were injected into the knee joints of young mice. One week after the injection, DMM surgery was conducted to induce OA. Since we expected that Gata4 overexpression would accelerate OA progression, we harvested samples for analysis 6 weeks after DMM.

**Figure 5:**
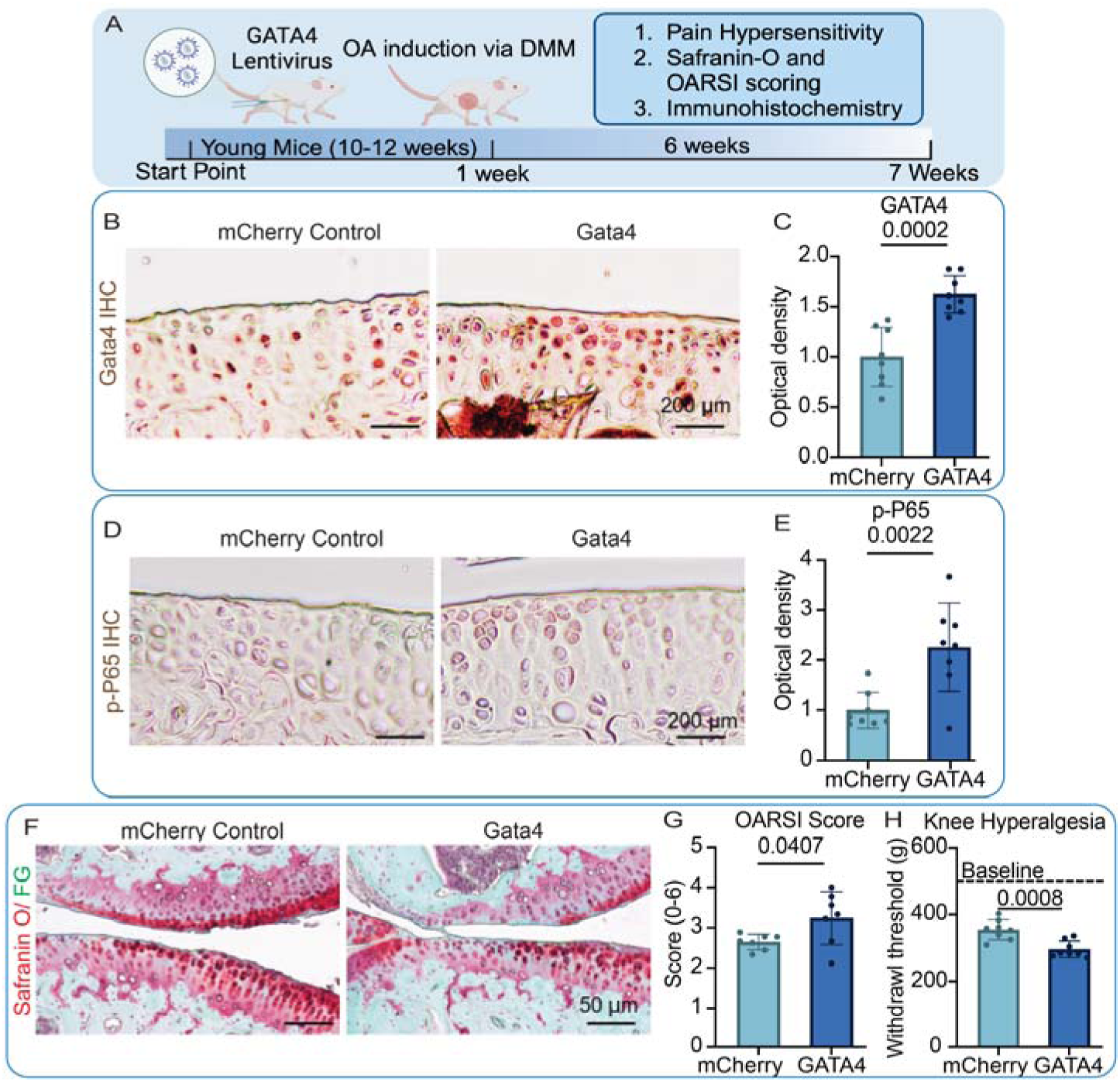
Gata4 overexpression in the knee joints accelerates OA progression in mice. (A) schematic of the study. Mice received one intraarticular injection of lentiviral vectors that carried mCherry or Gata4 gene one week before DMM surgery was performed. Knee joints were harvested 6 weeks post-surgery. Levels of Gata4 (B, C) and p-P65 (D, E) were assessed with IHC(B&D), and the staining was semi-quantitated with Image J (C&E). Cartilage degradation was assessed with (F) Safranin O/ fast green (FG) staining, and (G)OARSI score was calculated. (H)Knee hyperalgesia 6 weeks post-surgery. 507 g was the threshold baseline for non-surgery mice (dashed line). Student’s two-tailed t-test with Welch’s correction for standard deviation and a p-value of 0.05 was used for all statistical analysis. Created in BioRender. Makarczyk, M. (2025) https://BioRender.com/luon9t1

Results from IHC indicated a successful Gata4 overexpression in hyaline cartilage even 6 weeks after injection (**Figure 5B, C**). Interestingly, mice overexpressing Gata4 in the knee joint resulted in the elevation of p-P65 (**Figure 5D, E**), suggesting increased inflammation. Moreover, the mice in the Gata4 group displayed more severe OA and higher knee hyperalgesia than the control group, as revealed by a lower withdrawal threshold. (**Figure 5F-H**). Interestingly, mice from the Gata4 group displayed higher synovial inflammation (**Figure S8**).

## Discussion

The causal relationship between aging and OA has been documented, and understanding the molecular mechanisms is essential for the development of treatment methods. In this study, we discovered the critical roles of aging-associated increases in GATA4 levels. Specifically, overexpressing GATA4 in young chondrocytes impaired their capacity to form normal hyaline cartilage, while suppressing GATA4 in old chondrocytes restored their chondrogenic potential. We also demonstrated that GATA4 functions partially by promoting the activation of SMAD1/5. Our *in vivo* work further confirmed that high *Gata4* expression accelerated OA progression in mice. Lastly, we defined a small-molecule GATA4 inhibitor that can partially restore the capacity of old chondrocytes to create healthy cartilage, representing a potential DMOAD for further validation in the future.

Given the recognized challenges in harvesting healthy cartilage tissues from donors without arthritis, there are limited reports investigating chondrocyte aging per se. The current findings include proliferation and post-expansion chondrogenic capacity reduction with aging,^29^ increased MMP-13 production in response to catabolic stimuli,^30^ and altered response to TGF-β.^12^ One of our recent studies again demonstrated that aged chondrocytes displayed a reduced proliferation potential compared to young cells. Additionally, cartilage tissues generated by old chondrocytes contained more senescent cells than those from young cells.^31^ However, to the best of our knowledge, there were no publications describing the transcriptomic comparisons between young and old chondrocytes, which would be informative in defining targets to stop or reverse chondrocyte aging.

Through transcriptomic analysis, we were able to assess the expression of different genes and genetic pathways that occur as chondrocytes age. Of note, Hypoxia-Inducible Factor 1α (HIF1α) was the most differentially expressed gene predicted to regulate chondrocyte aging. The connection between HIF1α and aging has been previously reported.^32^ Furthermore, additional studies have investigated HIF1α in association with OA and assessed its use as a therapeutic target.^33,34^ Therefore, we decided to focus on GATA4, which was less studied in chondrocytes but highly associated with cellular senescence, an aging hallmark. However, our selection did not dampen the importance of HIF1α and other molecules listed in Figure 1D in chondrocyte aging. They can be further studied in the future using the same strategy employed in the current work.

The GATA family consists of several proteins (GATA1-6) with different variations of DNA-binding domains composed of zinc finger structures.^25^ Particularly, GATA4, 5, and 6 are involved in the development of the mesoderm and endoderm tissues,^35^ in which GATA4 has been detected in the development of structures such as the heart,^36,37^ pancreas,^38^ lung, and liver.^39,40^ In addition, GATA4 and GATA6 are the only members that are associated with aging.^41^ Particularly, GATA4 is unique in that it regulates tissues in a context-dependent manner and adopts a multifaceted role in the body, contributing to other age-related diseases such as atherosclerosis^42^ and heart failure.^43^

Furthermore, GATA4 might be associated with metabolic regulation. A study conducted by Patankar et al. investigated how GATA4 regulates obesity. Specifically, they used intestine-specific *Gata4* knockout mice to study diet-induced obesity, showing that the knockout mice were resistant to the high-fat diet, and that glucagon-like peptide-1 (GLP-1) release was increased. These findings indicated a decreased risk for the development for insulin resistance in knockout mice.^44^ This work was taken a step further in a subsequent publication, in which the same team investigated the dietary lipid-dependent and independent effects on the development of steatosis and fibrosis in *Gata4* knockout mice. The results from this work suggested that the knockdown of *Gata4* increases GLP-1 release, in turn suppressing the development of hepatic steatosis and fibrosis ultimately blocking hepatic *de novo* lipogenesis.^45^ These studies are especially interesting with the rise of GLP-1 based therapy for the treatment of OA.^46,47^ Thus, the coupling of GATA4-related metabolic dysfunction and OA should be further investigated.

In 2015, GATA4 was first introduced as a senescence regulator in a study conducted by Kang et al.^13^ This work demonstrated the role of GATA4 in human fibroblasts and determined that the inhibition of autophagy caused GATA4 accumulation following DNA damage. They examined the presence of GATA4 in the human brain and found that there was an increase of GATA4 in the prefrontal cortex of aged human samples.^13^ Further studies have shown that GATA4 regulates angiogenesis and inflammation in fibroblast-like synoviocytes in rheumatoid arthritis, indicating that GATA4 is required for the inflammation induced by IL-1β. This study also demonstrated that GATA4 binds to promoter regions on Vascular Endothelial Growth Factor (VEGF)-A and VEGFC to enhance transcription and regulate angiogenesis.^15^ In chondrocytes specifically, previous studies demonstrated that increased GATA4 levels are associated with chondrocyte senescence,^17,48,49^ an important change often observed in OA chondrocytes. Moreover, suppressing GATA4 with siRNA was shown to effectively abolish ionizing radiation-induced senescent phenotype in chondrocytes.^17^ Of note, all these studies relevant to chondrocytes were conducted *in vitro.* Herein, we, for the first time, demonstrate the role of GATA4 in regulating chondrocyte aging.

A theory of chondrocyte aging proposed by van der Kramm suggests that alterations to the TGF-β pathway induce chondrocyte hypertrophy and result in articular cartilage that is prone to OA development.^12^ While investigating the function of GATA4 in chondrocytes, we assessed how its levels contribute to TGF-β alterations in chondrocytes and found that GATA4 levels negatively correlated with the anabolic potential of chondrocytes (**Figures 2 and 4**). Our study indicated that there was an observed decrease in chondrogenesis and an increase in hypertrophy-related genes following *GATA4* overexpression (**Figure 2G)**. To maintain healthy cartilage homeostasis, numerous pathways are involved. In particular, TGF-β is a crucial cytokine necessary for cartilage homeostasis during OA,^50,51^ and aged chondrocytes respond differently to TGF-β compared to their young counterparts.^52^

Mechanistically, in the TGF-β pathway, TGF-β binds to the heterotetrameric receptor complex, which can be grouped into three receptor types (type I, type II, and type III). When TGF-β binds to its corresponding receptor, the activin-receptor-like kinases (ALKs) are activated.^53^ Typically, the anabolic binding of TGF-β to ALK4/5 results in the phosphorylation of SMAD2/3, protecting chondrocytes from hypertrophy.^50^ However, there can also be catabolic effects associated with the expression of SMAD1/5, typically expressed when TGF-β binds to ALK1/2/3/6.^50,53^ The phosphorylation of SMAD1/5 results in the promotion of ECM degrading proteins such MMP-13.^28,50^ As mentioned previously, research has demonstrated that aged chondrocytes respond differently to TGF-β compared to young chondrocytes.^52,54^ Chondrocyte aging has been linked to the increase of pSMAD1/5 signaling.^28^ These previous studies and literature review inspired us to explore the potential association between GATA4 levels and the activation of SMAD1/5.

Our results found that GATA4 resulted in the phosphorylation of SMAD1/5 even without TGF-β stimulation, which however did not alter the phosphorylation of SMAD2/3. Although there are no current publications specifying the complex relationship of GATA4 and SMAD1/5 in chondrocytes, a prior study reported that GATA4 was regulated by SMAD1/5. Specifically, SMAD1/5 and GATA4 can bind together to promote IL-6 expression in macrophages.^55^ In this study, it was shown that GATA4 was necessary for bone morphogenic protein-6 (BMP-6) mediated IL-6 induction, in which there are multiple GATA binding domains on the IL-6 promoter. This work further showed that GATA4 interacts with SMAD 2,3 and 4.^55^ Studies have suggested that BMP pathways and GATA4 work synergistically to regulate SMAD signaling.^56^ This information indicates that the involvement of GATA4 in the TGF-β signaling pathway is complex, and further studies should be conducted to better assess this relationship.

Additionally, a common hallmark of chondrocyte aging is the alternation of ECM, including composition change^2^ and stiffening.^57^ The mechanical integrity of ECM can directly affect chondrocyte phenotype and proliferation, and contribute to OA.^58^ For example, aging induces increased lysyl oxidase (LOX)-mediated collagen cross-linking and non-enzymatic advances glycation end-products (AGEs) begin to accumulate, which stiffens the ECM collagen fibrils.^59,60^ On the other hand, aging is also associated with increase in aggrecan fragmentation and decrease in the GAG length and packing density,^61^ which leads to decreased osmotic pressure and compressive resistance. It was found that the indentation modulus of murine cartilage decreases with age, likely associated with the reduced aggrecan content, integrity and osmotic pressure-endowed pre-tension within collagen fibrils.^60^ Meanwhile, earlier studies also indicate a softening effect of aging on cartilage tissue modulus.^62,63^ To this day, there is still a paucity of knowledge on the mechanism by which aging impacts the ECM and its reciprocal interplay with resident cells.

Investigating ECM alterations in conjunction with cellular senescence and TGF-β signaling could provide further insights into cartilage aging. A recent study by Fu et al. associated matrix stiffening with the promotion of chondrocyte senescence.^64^ Furthermore, matrix stiffening has been associated with modulating the TGF-β signaling pathway.^65–67^ Future studies should investigate the potential of matrix stiffening and the effect of GATA4 on pericellular matrix proteins such as decorin^68,69^, biglycan, collagen VI and XV, as these proteins assist with the regulation of biochemical interactions and assist with the maintenance of the chondrocyte microenvironment.^70^ Herein, the TGF-β signaling pathway can further alter the extracellular microenvironment^67^, which could promote cellular senescence and subsequently NF-κB pathway activation. Further investigation of ECM alterations and aging could elucidate the molecular events governing these changes, providing key insights on the interplay between chondrocytes and their environment in the context of aging.

While the TGF-β pathway is closely associated with cartilage matrix remodeling, we also found increases in the levels of multiple proinflammatory cytokines after GATA4 overexpression. We investigated the NF-κB pathway since it is also activated in aged tissues.^71,72^ During the aging process, NF-κB is thought to amplify and potentially propagate cellular senescence through the senescence-associated secretory phenotype (SASP).^16^ A study by Qiao et al. indicated that GATA4 regulates NF-κB in dental pulp cells^14^ and fibroblasts^15^. Specifically, using siRNAs, they determined that the knockdown of GATA4 decreased p65 production induced by lipopolysaccharide (LPS).^14^ Other studies have investigated the role of GATA4 in the synovium and the progression of the disease state of rheumatoid arthritis and OA. Jia et al. showed that the knockdown of GATA4 attenuated synovial inflammation and joint damage in a collagenase-induced arthritis mouse model.^15^ More recently, a study conducted by Chen et al. investigated the roles of GATA4 in fibroblast-like synoviocytes (FLS) and determined that GATA4 induced cellular senescence in FLSs in OA progression.^27^ Our study also discovered that the siRNA knockdown of GATA4 decreased the phosphorylation of p65 in aged chondrocytes (**Figure 4**). However, we also noticed that MMP-13 expression levels decreased in both *GATA4* overexpression and knockdown experiments. It should be noted that MMP-13 is constitutively produced in human chondrocytes but is only activated under pathological conditions. Given that MMP-13 is regulated by different transcriptional factors and cytokines, as well as RNAs,^28^ its associations with GATA4 require further investigation in the future. Collectively, these findings further indicate that GATA4 might regulate aging partially through TGF-β and NF-κB pathways.

Our current study has not fully explored why aging promotes GATA4 expression. The study from Kang et al. indicated that DNA damage contributed to GATA4 accumulation.^13^ It is known that DNA damage response (DDR) is regulated by ataxia telangiectasia mutated (ATM) and ataxia telangiectasia and Rad3–related (ATR) signaling.^13,73,74^ The activation of these signaling pathways inhibits autophagy-related protein p62.^75^ In addition, Copp et al. reported increased DNA damage in old chondrocytes.^5^ These studies implied that DNA damage may be a reason for GATA4 upregulation. Our preliminary data and prior work from Kan et al.^13^ support this possibility. Specifically, a DNA damage-inducing agent, doxorubicin, promoted the upregulation of GATA4 in chondrocytes (**Supplementary Figure S9**). To further link DNA damage to GATA4 accumulation, a study by Chung et al. used Tributyltin (TBT), a well-known endocrine-disrupting chemical, to induce DNA damage in articular chondrocytes. After 24 hours of TBT treatment, there was a significant increase in *GATA4* expression and expression of senescence markers.^48^ The study by Kang et al. demonstrated that the suppression of p62 following DNA damage leads to GATA4 accumulation due to the lack of autophagy.^13^ DNA damage is known to increase with age.^76^ Therefore, we believe that DNA damage due to aging is a key driver of the upregulation of GATA4 in old chondrocytes.

In conclusion, we have determined that GATA4 is increased in aged chondrocytes compared to young in both humans and mice, which may be induced by increased DNA damage observed in aged cells. We also demonstrated that GATA4 overexpression impaired the quality and quantity of cartilage created by chondrocytes and accelerated OA progression in mice. Conversely, suppressing GATA4 with siRNA or small-molecule inhibitors partially restores the capacity of old chondrocytes to form cartilage. Additionally, our study found that GATA4 can activate SMAD1/5 and change chondrocyte response to TGF-β. Overall, our study indicated that GATA4 could be a contributor to OA onset and progression in aged individuals, which can also serve as a potential target to prevent aging-associated OA.

There are some limitations of this study that can be further addressed in future studies. First, this study has allowed us to examine the potential mechanisms of GATA4 in chondrocyte aging. Although we found that GATA4 was generally increased with aging, some young donors also exhibited increased levels of GATA4, which may be associated with increased DNA damage, as discussed above, or other stressors. Therefore, GATA4 should be used together in conjunction with other aging biomarkers, such as epigenetic clock ^77^ to precisely define chondrocyte aging. Future work should examine biological versus chronological aging and epigenetic clock-based assessments to explain the variabilities in GATA4 expression among donors. Second, the TGF-β signaling pathway is complex, and there are multiple studies that have associated it with GATA4. However, the relationship between GATA4 overexpression and SAMD1/5 activation should be further investigated to elucidate specific signaling mechanisms. Third, during our *in vivo* work, the intraarticular injection of GATA4 lentivirus was not chondrocyte-specific. Therefore, the injection also allowed for other cell types to overexpress GATA4. Future work should be conducted using transgenic mouse lines for cartilage-specific inducible overexpression or depletion of *Gata4* to further investigate the role of GATA4 in chondrocytes. Furthermore, studies using GATA4 knockdown do not demonstrate a complete reversal of aging in chondrocytes, and additional *in vivo* assessment needs to be conducted to verify whether GATA4 could be a therapeutic target for chondrocyte aging. In particular, our *in vitro* study demonstrated the potential of using small-molecule GATA4 to enhance the quality of cartilage created by old chondrocytes. We can validate the findings *in vivo*, as well as develop other GATA4 inhibitors. Lastly, our work indicates that GATA4 has a role in chondrocyte aging, but that does not negate the other aging factors and pathways. Combining current findings and investigating the prevalence of GATA4 in association with other aging molecules will further contribute to our understanding of the role of GATA4 in aging and possible OA onset.

## Acknowledgments

This work was supported by the Department of Orthopaedic Surgery and the Orland Bethel Family Musculoskeletal Research Center (BMRC) at the University of Pittsburgh, as well as the University of Pittsburgh Center for Research Computing through the resources provided. This study used the Luminex Core Facility of UPCI. M.J.M was supported by the NIH T32 EB001026-Cellular Approaches to Tissue Engineering and Regeneration, Cardiovascular Bioengineering Training Program T32-HL076124, and Ruth L. Kirschstein National Research Service Award (NRSA) Individual Predoctoral Fellowship (Parent F31) 1F31AR083814 - 01A1. The authors thank the funding support from the NIH (P30AG038072). The Chicago Center on Musculoskeletal Pain Research Core Center, supported by NIH P30AR079206, provided training on pain-assessing methods. Schematic figures were created at https://BioRender.com.

## Data Availability

The Raw and processed RNA-seq data was uploaded to Gene Expression Omnibus (GEO) with accession ID: GSE287861. The data will be publicly available when the manuscript is accepted.

## Notes

### Competing Interest Statement

The authors have declared no competing interest.

### Summary of Updates

We discussed the changes in the ECM during the aging process. We also added more justification for selecting GATA4.

