## Supplemental materials for "Aging-associated Increase of GATA4 levels in Articular Cartilage is Linked to Impaired Regenerative Capacity of Chondrocytes and Osteoarthritis"

**Supplementary Figures and Tables**

**Figure S1**

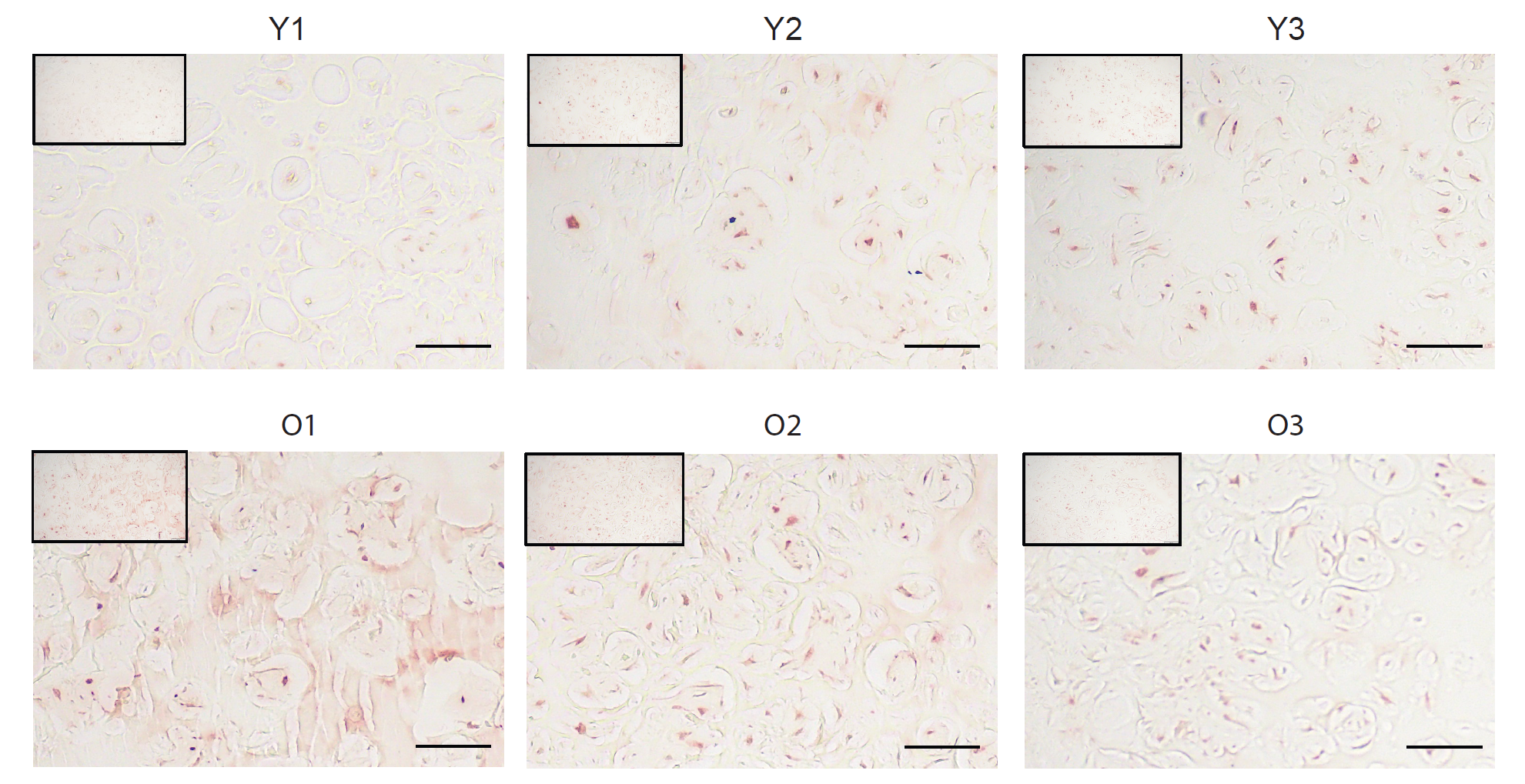

***Figure S1.*** *GATA4 IHC of healthy human cartilage tissue from 3 young (Y1,Y2, Y3) and aged (O1,O2, O3) donors. Scale Bar=50 µm.*

**Figure S2**

Young	old

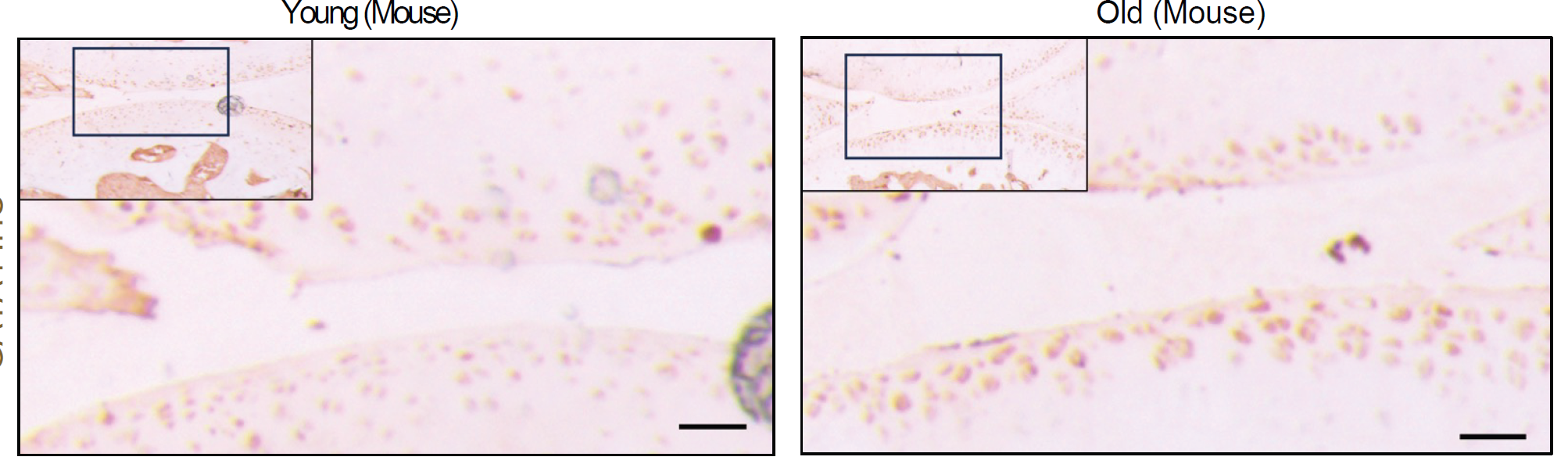

***Figure S2.*** *GATA4 IHC of healthy mouse cartilage tissue from young (left) and old (right) mice. Scale Bar=50 µm.*

**Figure S3**

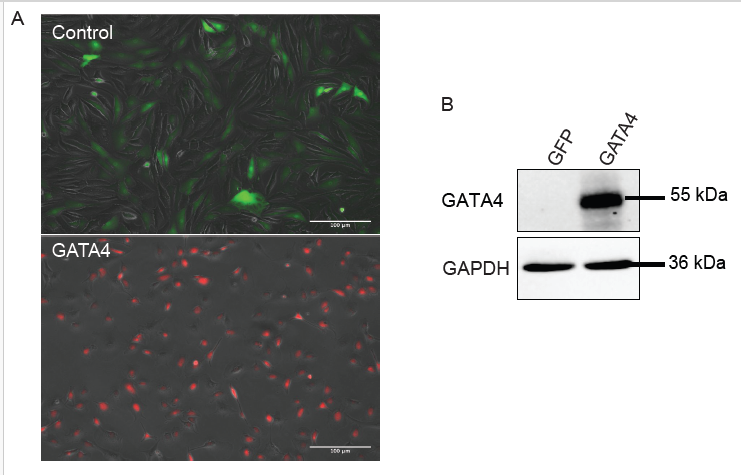

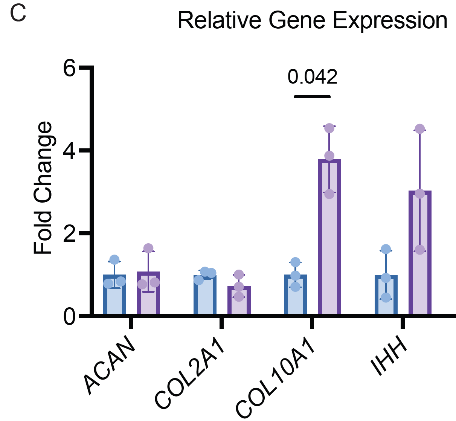

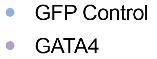

***Figure S3.*** *(A) GATA4 overexpression of young, pooled chondrocytes in monolayer culture 48 hours after infection. The lentiviral control contained the EF1A promoter-driven expression of Green Fluorescent Protein (GFP), and the GATA4 lentivirus contained the EF1A promoter-driven overexpression of GATA4 with dTomato fluorescent protein. Scale Bar=100 µm. (B) Western blot to measure GATA4 protein levels. (C) RT-qPCR analysis of relative gene expression of cartilage matrix proteins aggrecan (ACAN) and collagen type II-α1 (COL2A1), and hypertrophy markers collagen type X- α1 (COL10A1) and India hedgehog (IHH).*

**Figure S4**

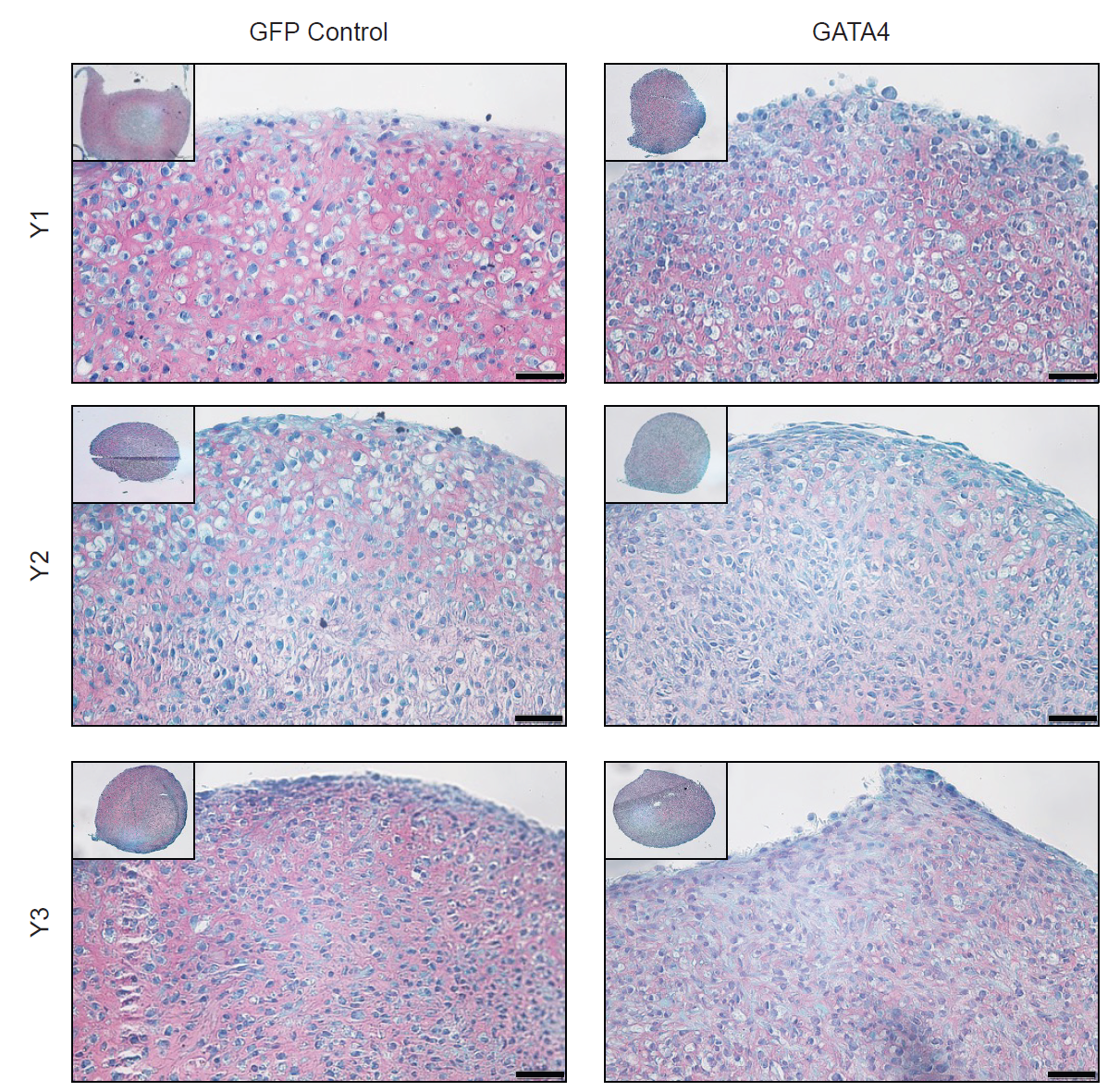

***Figure S4.*** *Safranin-O staining for pellets derived from young, individual chondrocyte (Y1-3) transduced with GFP control lentivirus or GATA4 lentivirus. Pellets were cultured in chondrogenic medium for 7 days. Scale Bar=50 μm.*

***Figure S5***

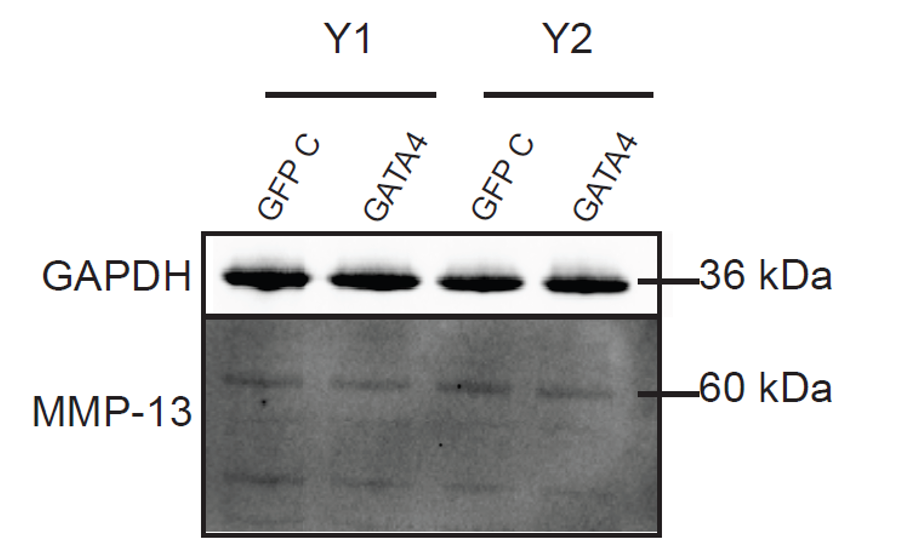

***Figure S5.*** *Western blot to examine MMP-13 protein levels in pellets derived from two young chondrocyte lines (Y1 and Y2) overexpressing GFP control (GFP C) or GATA4.*

***Figure S6***

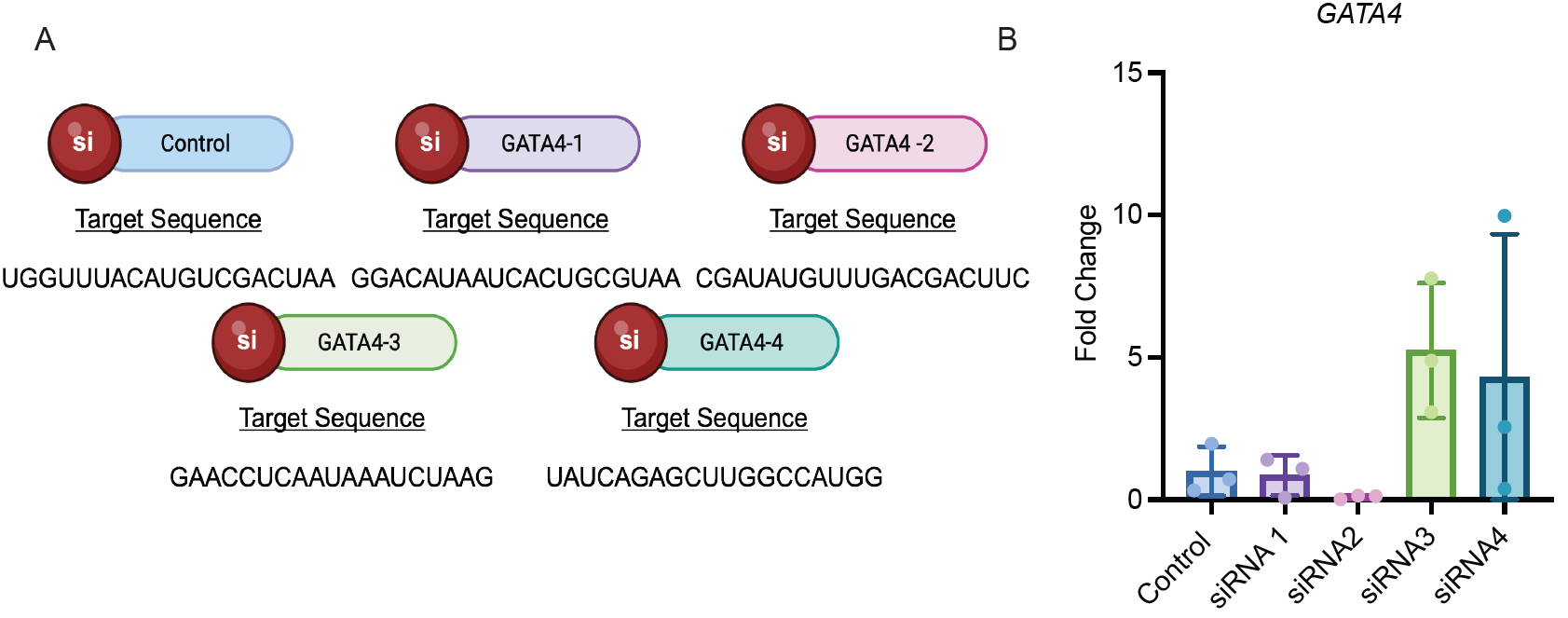

***Supplementary Figure 6. Assessment of GATA4 siRNAs in monolayer****. (A) Schematic of the four different GATA4 siRNAs with corresponding target sequences assessed for GATA4 knockdown in monolayer culture of old pooled chondrocytes. (B) RT-qPCR assessing GATA4 levels after siRNA treatment (n=3). Created in BioRender. Makarczyk, M. (2025) https://BioRender.com/luon9t1*

**Figure S7**

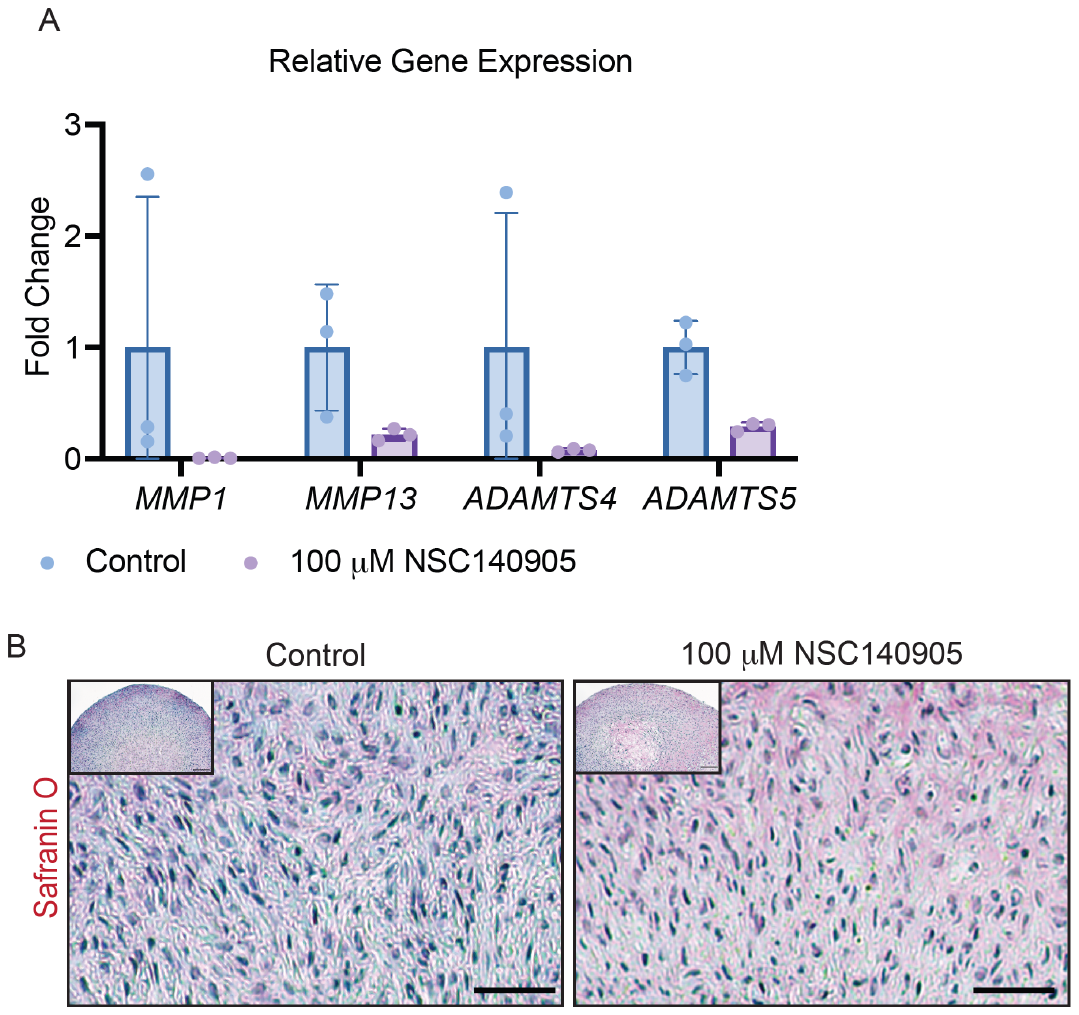

***Supplementary Figure 7. Assessment of GATA4 small molecule inhibitor,*** ***NSC140905.*** *Pooled old human chondrocytes were pelleted and treated with the chondrogenic medium with or without supplanting NSC140905 for 14 days.* *(A) RT-qPCR analysis of relative gene expression of matrix-degrading enzymes, including MMP-1,13 and ADAMTS 4 and 5 (n=3). Student’s two-tailed t-test with Welch’s correction for standard deviation and a p-value of 0.05 was used for all statistical analysis. (B) Safranin-O/Fast green staining. Bar=50 μM.*

**Figure S8**

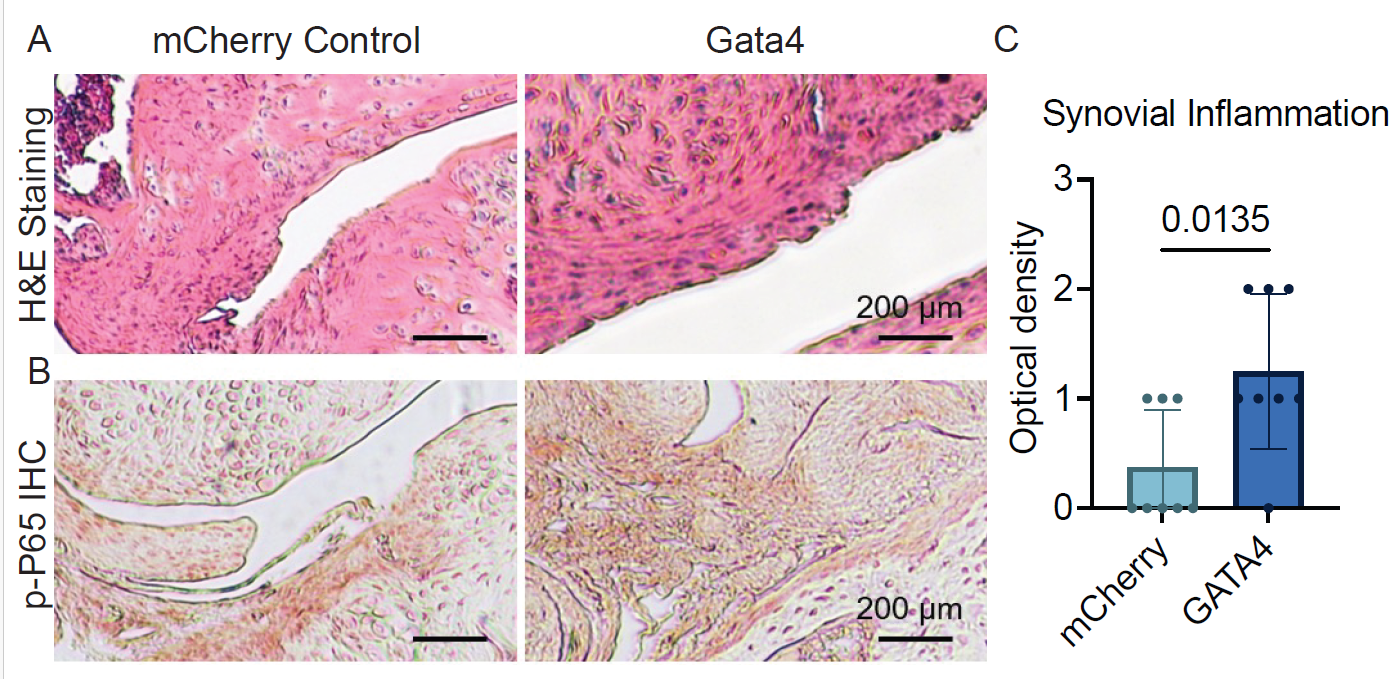

***Supplementary* *Figure S8.*** *(A) H&E staining and (B) p-P65 IHC to assess synovial inflammation in mice treated with lentiviral vectors carrying mCherry Control or Gata4.* C. p-P65 IHC staining was also semi-quantitated.

**Figure S9**

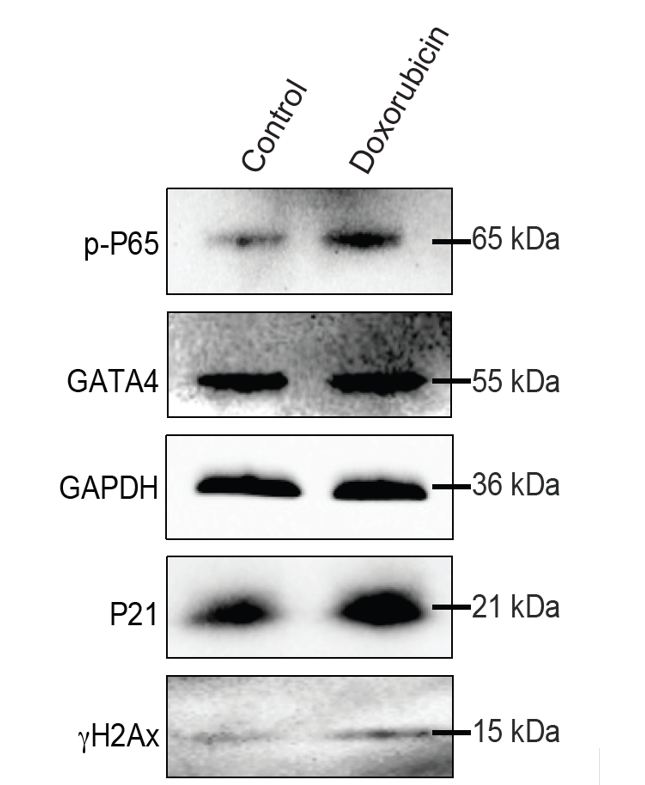

***Supplementary* *Figure S9.*** *Wester blot to examine protein levels in chondrocytes treated with doxorubicin (100nM) or vehicle control for 3 days.*

**Table S1. Information of chondrocyte donors**

RNA sequencing

| **Age** | **Gender** |
| --- | --- |
| 24 | Female |
| 36 | Female |
| 38 | Female |
| 70 | Female |
| 73 | Female |
| 74 | Female |

Western Blot

| **Age** | **Gender** |
| --- | --- |
| 22 | Male |
| 26 | Male |
| 29 | Male |
| 69 | Male |
| 70 | Male |
| 73 | Female |

Human Cartilage Tissue IHC

| **Age** | **Gender** |
| --- | --- |
| 24 | Male |
| 25 | Female |
| 27 | Female |
| 33 | Male |
| 67 | Female |
| 76 | Female |
| 78-1 | Female |
| 78-2 | Male |

Young Chondrocyte Pool

| **Age** | **Gender** |
| --- | --- |
| 22 | Male |
| 29 | Male |
| 38 | Female |
| 42 | Female |

Old Chondrocyte Pool #1

| **Age** | **Gender** |
| --- | --- |
| 66 | Male |
| 70 | Male |
| 73 | Female |

Old Chondrocyte Pool #2

| **Age** | **Gender** |
| --- | --- |
| 66-1 | Male |
| 66-2 | Male |
| 70-1 | Male |
| 70-2 | Male |

Monolayer young individual chondrocytes

| **Age** | **Gender** |
| --- | --- |
| 29 | Male |
| 36 | Female |
| 38 | Female |

Pellet young individual chondrocytes

| **Age** | **Gender** |
| --- | --- |
| 21 | Male |
| 35 | Male |
| 38 | Female |

**Table S2. Antibodies used for immunofluorescence (IF), Immunohistochemistry (IHC), or Western blot (WB)**

| **Antibody** | **Origin** | **Cat. No** | **Species** | **Assay** | **Dilution** |
| --- | --- | --- | --- | --- | --- |
| Anti-Human Collagen Type II | MP Biomedicals | SKU 0863171 | Mouse | IHC | 1:200 |
| Biotinylated Goat Anti-Rabbit IgG | Vector Laboratories | PK-6101 | Rabbit | IHC | 1:50 |
| Goat Anti-Rabbit IgG H&L (HRP) | Abcam | Ab6721 | Goat | WB | 1:5000 |
| GAPDH (D16H11) XP® Rabbit mAb | Cell Signaling Technology | 5174s | Rabbit | WB | 1:2000 |
| Smad1 (D59D7) XP® Rabbit mAb | Cell Signaling Technology | 6944s | Rabbit | WB | 1:500 |
| Anti-GATA4 antibody (ab84593)- Discontinued | Abcam | ab84593 | Rabbit | IHC/WB | 1:500 |
| GATA-4 (D3A3M) Rabbit mAb | Cell Signaling Technology | 369366s | Rabbit | IHC/WB | 1:1000 |
| Phospho-Smad1/5 (Ser463/465) (41D10) Rabbit mAb | Cell Signaling Technology | 9516s | Rabbit | WB | 1:500 |
| Phospho-Smad2 (Ser465/467) (138D4) Rabbit mAb | Cell Signaling Technology | 3108s | Rabbit | WB | 1:500 |
| Smad2/3 (D7G7) XP® Rabbit mAb | Cell Signaling Technology | 8685s | Rabbit | WB | 1:500 |
| Anti-NF-kB p65 (phospho S536) antibody [EP2294Y] | Abcam | ab76302 | Rabbit | WB | 1:500 |
| Anti-NF-kB p65 antibody [E379] | Abcam | ab32536 | Rabbit | WB | 1:500 |
| Anti-p21 antibody [EPR362] - BSA and Azide free | Abcam | ab218311 | Rabbit | WB | 1:1000 |
| Phospho-Histone H2A.X (Ser139) (20E3) Rabbit mAb | Cell Signaling Technology | 9718s | Rabbit | WB | 1:500 |

**Table S3: Full names of genes shown in Figure 1**

| Transcription Regulator Name | Acronym |
| --- | --- |
| Hypoxia-inducible factor 1-alpha | HIF1A |
| GATA binding protein 4 | GATA4 |
| MAF bZIP transcription factor B | MAFB |
| Homeobox D10 | HOXD10 |
| CCAAT enhancer binding protein alpha | CEBPA |
| Forkhead box L2 | FOXL2 |
| Caudal type homeobox 2 | CDX2 |
| Early Growth Response 1 | EGR1 |
| Signal Transducer and Activator of Transcription 3 | STAT3 |
| Paired Box 1 | PAX1 |
| Sequestosome 1 | SQSTM1 |
| Myocardin Related Transcription Factor B | MRTFB |
| WW Domain Binding Protein 2 | WBP2 |
| Scleraxis bHLH Transcription Factor | SCX |
| Twist Family bHLH Transcription Factor 1 | TWIST1 |
| Myocyte Enhancer Factor 2D | MEF2D |
| Lysine Methyltransferase 2D | KMT2D |
| Achaete-scute Family bHLH Transcription Factor 1 | ASCL1 |
| Recombination Signal Binding Protein for Immunoglobulin kappa J region | RBPJ |
| SIX homeobox 1 | SIX1 |
| FERM domain containing 3 | FRMPD3 |
| Ceruloplasmin | CP |
| SPARC related modular calcium binding 2 | SMOC2 |
| ATP binding cassette subfamily A member 13 | ABCA13 |
| Inositol 1,4,5-triphosphate receptor type 1 | ITPR1 |
| NIM1 serine/threonine protein kinase | NIM1K |
| Fibroblast growth factor 13 | FGF13 |
| Ankyrin repeat domain 12 | ANKRD12 |
| GTP binding protein overexpressed in skeletal muscle | GEM |
| erythroferrone | FAM132B |
| Pappalysin 1 | PAPPA |
| PPARG coactivator 1 alpha | PPARGC1A |
| Synaptotagmin like 2 | SYTL2 |
| Leucine rich repeat transmembrane neuronal 2 | LRRTM2 |
| G protein-coupled receptor class C group 5 member B | GPRC5B |
| Polypeptide N-acetylgalactosaminyltransferase 15 | GALNT15 |
| Growth associated protein 43 | GAP43 |
| Fibroblast growth factor binding protein 1 | FGFBP1 |
| ATRX chromatin remodeler | ATRX |
| Protocadherin gamma subfamily B, 2 | PCDHGB2 |
| Calsyntenin 2 | CLSTN2 |
| Fibroblast growth factor binding protein 2 | FGFBP2 |
| Desmocollin 2 | DSC2 |
| TNF receptor superfamily member 21 | TNFRSF21 |
| Kinesin family member 13B | KIF13B |
| Plexin A2 | PLXNA2 |
| Cysteine rich protein 2 | CRIP2 |
| Cysteine rich protein 1 | CRIP1 |
| KLF transcription factor 2 | KLF2 |
| Serum/glucocorticoid regulated kinase 1 | SGK1 |
| LIM zinc finger domain containing 2 | LIMS2 |
| Long intergenic non-protein coding RNA 1133 | LINC01133 |
| Adipogenesis regulatory factor | ADIRF |
| Ornithine decarboxylase 1 | ODC1 |
| Kazrin, periplakin interacting protein | KAZN |
| Chondroitin polymerizing factor | CHPF |
| RAB23, member RAS oncogene family | RAB23 |
| Alkaline phosphatase, biomineralization associated | ALPL |
| Signal transducer and activator of transcription 4 | STAT4 |
| G protein-coupled receptor 1 | GPR1 |
| Spectrin alpha, non-erythrocytic 1 | SPTAN1 |
| Very low density lipoprotein receptor | VLDLR |
| Fatty acid desaturase 3 | FADS3 |
| Cancer susceptibility candidate 4 | CASC4 |
| NAD(P)H quinone dehydrogenase 1 | NQO1 |
| Pyrroline-5-carboxylate reductase 1 | PYCR1 |
| ERBB receptor feedback inhibitor 1 | ERRFI1 |
| NmrA-like family domain containing 1 pseudogene | LOC344887 |
| Desumoylating isopeptidase 2 | DESI2 |
| Basonuclin zinc finger protein 1 | BNC1 |

**Table S4. Primers for qRT-PCR**

| **Gene** | **Forward primer (5’-3’)** | **Reverse primer (5’-3’)** |
| --- | --- | --- |
| *RPL13A* | GCCATCGTGGCTAAACAGGTA | GTTGGTGTTCATCCGCTTGC |
| *GATA4* | CAGTCTACGTGCCCACACC | TCCCGCCTGGCTCCAT |
| *ACAN* | AGTCACACCTGAGCAGCATC | AGTTCTCAAATTGCATGGGGTGTC |
| *COL2A1* | GGATGGCTGCACGAAACATACCGG | CAAGAAGCAGACCGGCCCTATG |
| *COL10A1* | CCCTCTTGTTAGTGCCAACC | AGATTCCAGTCCTTGGGTCA |
| *IHH* | AACTCGCTGGCTATCTCGGT | GCCCTCATAATGCAGGGACT |
| *IL-6* | ACTCACCTCTTCAGAACGAATTG | CCATCTTTGGAAGGTTCAGGTTG |
| *IL-8* | TTTTGCCAAGGAGTGCTAAAGA | AACCCTCTGCACCCAGTTTTC |
| *TNF-α* | CCTCTCTCTAATCAGCCCTCTG | GAGGACCTGGGAGTAGATGAG |
| *MMP-1* | AAAATTACACGCCAGATTTGCC | GGTGTGACATTACTCCAGAGTTG |
| *MMP-2* | GGTCACATCGCTCCAGACT | TACAGGATCATTGGCTACACACC |
| *MMP-3* | CGGTTCCGCCTGTCTCAAG | CGCCAAAAGTGCCTGTCTT |
| *MMP-12* | GGAATCCTAGCCCATGCTTTT | CATTACGGCCTTTGGATCACT |
| *MMP-13* | ACTGAGAGGCTCCGAGAAATG | GAACCCCGCATCTTGGCTT |
| *ADAMTS4* | GAGGAGGAGATCGTGTTTCCA | CCAGCTCTAGTAGCAGCGTC |
| *ADAMTS5* | GAACATCGACCAACTCTACTCCG | CAATGCCCACCGAACCATCT |

**Table S5. Information of Luminex assay kit**

| **Target Molecule** | **Method** | **Catalog number & supplier** |
| --- | --- | --- |
| IL-8 | Luminex® assay | HAGP1MAG-12K, EMD Millipore |
| IL-6, CCL2, MMP-1, MMP-2, MMP-13. | Luminex® assay | LXSAHM-18, R&D Systems |

**Supplementary Table 6: Comprehensive list of proteins assessed in LUMINEX**

| **Target Molecule** | Catalog number & supplier |
| --- | --- |
| Interleukin-6 (IL-6) | LXSAHM-18, R&D Systems |
| Interleukin-13 (IL-13) | LXSAHM-18, R&D Systems |
| Matrix Metallopeptidase-3 (MMP-3) | LXSAHM-18, R&D Systems |
| Periostin (OSF-2) | LXSAHM-18, R&D Systems |
| Vascular Endothelial Growth Factor Receptor 2 (VEGFR2) | LXSAHM-18, R&D Systems |
| MMP-8 | LXSAHM-18, R&D Systems |
| Adiponectin (AdipoQ) | LXSAHM-18, R&D Systems |
| Complement Factor D/Adipsin (CFD) | LXSAHM-18, R&D Systems |
| Ectonucleotide Pyrophosphatase/Phosphodiesterase 2 Autotaxin (ENPP-2) | LXSAHM-18, R&D Systems |
| MMP-2 | LXSAHM-18, R&D Systems |
| Osteopontin (OPN) | LXSAHM-18, R&D Systems |
| C-C motif ligand 2 (CCL2) | LXSAHM-18, R&D Systems |
| (C-X-C motif) ligand 1 (CXCL1) | LXSAHM-18, R&D Systems |
| IL-1ra | LXSAHM-18, R&D Systems |
| MMP-1 | LXSAHM-18, R&D Systems |
| MMP-13 | LXSAHM-18, R&D Systems |
| Tissue Inhibitor of Metalloproteinase-1 (TIMP-1) | HTMP1MAG-54K, EMD Millipore |
| TIMP-2 | HTMP1MAG-54K, EMD Millipore |
| Angiopoietin-2 (Ang-2) | HAGP1MAG-12K, EMD Millipore |
| Bone Morphogenic Protein-9 (BMP-9) | HAGP1MAG-12K, EMD Millipore |
| Epidermal Growth Factor (EGF) | HAGP1MAG-12K, EMD Millipore |
| Endoglin (CD105) | HAGP1MAG-12K, EMD Millipore |
| Endothelin-1 (ET-1) | HAGP1MAG-12K, EMD Millipore |
| Fibroblast Growth Factor-1 (FGF-1) | HAGP1MAG-12K, EMD Millipore |
| FGF-2 | HAGP1MAG-12K, EMD Millipore |
| Follistatin (FSH) | HAGP1MAG-12K, EMD Millipore |
| Granulocyte Colony Stimulating Factor (G-CSF) | HAGP1MAG-12K, EMD Millipore |
| Heparin-Binding EGF-like Growth Factor HB-EGF | HAGP1MAG-12K, EMD Millipore |
| Hepatocyte Growth Factor (HGF) | HAGP1MAG-12K, EMD Millipore |
| IL-8 | HAGP1MAG-12K, EMD Millipore |
| Leptin (LEP) | HAGP1MAG-12K, EMD Millipore |
| Placental Growth Factor (PLGF) | HAGP1MAG-12K, EMD Millipore |
| VEGF-A | HAGP1MAG-12K, EMD Millipore |
| VEGF-D | HAGP1MAG-12K, EMD Millipore |
